# Cannabidiol dampens propagation of hippocampal hyperactivity and differentially modulates feedforward and feedback inhibition

**DOI:** 10.1101/2025.08.26.672420

**Authors:** Simon Chamberland, Evan C. Rosenberg, Erica R. Nebet, Orrin Devinsky, Richard W. Tsien

## Abstract

Cannabidiol (CBD) decreases seizures in patients with severe pediatric-onset epilepsies including Dravet, Lennox-Gastaut, and Tuberous Sclerosis syndromes. However, the effects of CBD on neuronal activity and circuits remain obscure. In the mouse hippocampus, we found that CBD causes a GPR55-independent decrease in CA1 pyramidal neuron firing frequency and a GPR55-dependent reduction in CA3 to CA1 hippocampal activity propagation. CBD-mediated decrease in high-frequency activity was mimicked by GPR55 antagonism and prevented by GPR55 deletion and blockade of GABAergic transmission. Dampening high-frequency activity was accompanied by increased recruitment of parvalbumin+ (PV)-INs and reduced recruitment of somatostatin+ (SST)-INs, leveraging the inhibitory subcircuit to limit propagation of hyperactivity. CBD-induced attenuation of high frequency spike propagation was mimicked by pharmacological enhancement and optogenetic engagement of PV-INs. Such increased on-demand recruitment of PV-INs dampened propagation of high-frequency activity to hippocampal CA1 similarly to CBD. We predict that CBD potentially curbs propagation and perpetuation of seizure activity via these mechanisms.

## Introduction

Temporal lobe epilepsy (TLE), the most prevalent epilepsy syndrome in humans, is caused by a pathological form of synchronous neuronal activity (Wiebe, 2000). The hippocampus and neighboring structures comprise temporal lobe neuronal circuits composed of abundant glutamatergic pyramidal neurons (PYRs) and limited GABAergic interneurons (INs). While less numerous, these INs are genetically and functionally diverse (Pelkey et al., 2017; Harris et al., 2018; Chamberland et al., 2024) and disproportionately impactful (Klausberger and Somogyi, 2008), operating in exquisite coordination to regulate PYR firing. In PYRs of hippocampal area CA1, distinct INs target multiple cellular domains: perisomatic inhibition curbs cell body depolarization and neuronal firing, while dendritic inhibition opposes synaptic integration and dendritic spikes (Miles et al., 1996; Vaasjo et al., 2025). Overall, inhibition serves as clockwork to drive ensembles of PYRs in rhythmic firing and hippocampal oscillations spanning a wide range of frequencies (Buzsaki, 2002; Klausberger and Somogyi, 2008).

Epileptic seizures are governed by the same circuit components operating in healthy brain but reflect excessive, potentially runaway activity. High-frequency oscillations underlie physiological information flow (Lisman, 1997; Abbott and Regehr, 2004; Izhikevich, 2007; Buzsaki and Silva, 2012; Buzsaki, 2015), but become pathological if neuronal coordination fails. IN dysfunction is a major cause of epileptiform activity (Prince et al., 2009; Liu et al., 2014; Paz and Huguenard, 2015; Buckmaster, Abrams and Wen, 2017). For example, Dravet Syndrome (DS), a devastating form of childhood-onset epilepsy, usually results from haploinsufficiency of Na_V_1.1 spike-generating sodium channels. In mice, DS is largely recapitulated by deleting these channels in parvalbumin (PV)-positive, fast-spiking interneurons (Cheah et al., 2012; Tai et al., 2014; but see Rubinstein et al., 2015).

Epilepsy therapies seek to downregulate principal neuron excitability while upregulating GABAergic feedforward inhibition (Paz and Huguenard, 2015). Sodium channel inhibition and positive allosteric modulation of GABA_A_Rs, key epilepsy therapies, leave a third of all epilepsy patients with ongoing seizures; i.e., treatment-resistant epilepsy (French and Gazzola, 2011; Kanner et al., 2018; Steriade and French, 2022; Scheffer et al., 2024). Inhibitory neurotransmission can protect circuits from pathological insults (Cossart et al., 2001) and modulation of INs activity directly impacts behavioral and electrographic seizures (Krook-Magnuson et al., 2013; Tung, Berglund and Gross, 2016; Christenson Wick and Krook-Magnuson, 2018; Levesque et al., 2023). Therapeutic agents selectively targeting INs could offer a potential well-tolerated epilepsy therapy.

*Cannabis sativa* has been used for centuries to treat epilepsy (Rosenberg et al., 2015), and provides a plethora of compounds with diverse pharmacological actions (Mechoulam, 2023; Devinsky et al., 2024). The non-euphoric phytocannabinoid cannabidiol (CBD) is effective in treating several forms of pediatric-onset epilepsies (Devinsky et al., 2017; Devinsky et al., 2018). The pharmacological profile of CBD is complex, directly modulating activity at many ion channels (Patel et al., 2016; Zhang et al., 2022; Ghovanloo, Arnold and Ruben, 2023) and G-protein coupled receptors, including GPR55 (Lauckner et al., 2008; Sylantyev et al., 2013; Kaplan et al., 2017; Rosenberg et al., 2023), and other targets (Ibeas Bih et al., 2015). These experimental findings offer diverse, yet potentially contradictory explanations for CBD’s clinical effectiveness, prompting investigation at the microcircuit level.

Here we focus on key unanswered questions encompassing excitatory and inhibitory evoked transmission. First, how does CBD modify circuit operations when a canonical hippocampal circuit is subjected to a physiological input pattern to drive hippocampal information transfer? Second, which CBD actions are GPR55-dependent, and which are not? Third, is CBD efficacy predominated by alteration of synaptic weights or of intrinsic membrane properties? Fourth, are CBD actions on inhibitory neurons and neurotransmission uniform across all IN subtypes, as expected for benzodiazepines, or specific to certain interneuron subtype(s), as observed for THC (Glickfeld and Scanziani, 2006) and in a different profile, CBD (Khan et al., 2018)? Delineation of basic mechanisms may improve our ability to use CBD as a therapeutic agent.

## Results

### Cannabidiol reduces the propagation of high-frequency activity from hippocampal CA3 to CA1

Unconstrained propagation of high-frequency activity from hippocampal CA3 to CA1 is a feature of epileptiform and ictal discharges in CA1 (Dzhala and Staley, 2003). CBD is an anti-seizure therapy in humans (Devinsky et al., 2017; Devinsky et al., 2018) and is anticonvulsant in multiple animal models (Rosenberg, Patra and Whalley, 2017; Patra et al., 2019; Devinsky et al., 2024). While CBD acts through multiple molecular mechanisms, its neuronal and circuit effects remain underexplored (Jones et al., 2010).

We examined how CBD (20 µM) modulates the propagation of high-frequency activity in acute hippocampal slices. For electrophysiological testing, we used a concentration of 20 μM, just below the Cmax attained in the brain after giving CBD 120 mg/kg (i.p.), near optimal anticonvulsant dosing (Jones et al., 2010). CA1 pyramidal neurons (CA1-PYRs) were recorded in current-clamp mode and Schaffer collaterals were electrically stimulated to generate action potentials (APs, 5 pulses delivered at 50Hz, Fig. 1A-B). The likelihood of synaptically-evoked APs was significantly reduced by CBD (Fig. 1A-B), while CBD had no effect on spontaneous firing or resting membrane potential of CA1-PYR (Fig. S1A-E). Therefore, CBD curtails the propagation of high-frequency activity. This dampening effect could result from modulation of synaptic inputs to CA1-PYRs or modification of the intrinsic excitability of CA1-PYRs, or both. We tested each possibility, first focusing on synaptic potentials evoked by stimulation of Schaffer collaterals. Decreasing stimulation intensity relative to experiments in Fig. 1A-B revealed large amplitude synaptic potentials that were significantly decreased by CBD (Fig. 1B-C). CBD binds to GPR55 and antagonizes the GPR55 ligand lysophosphatidylinositol (LPI) (Oka et al., 2007; Ryberg et al., 2007). Endogenous LPI is released during high-frequency activity and augments excitatory transmission (Sylantyev et al., 2013; Rosenberg et al., 2023). Consistent with this, bath application of LPI (4 µM) enhanced train-evoked PSPs transiently (p<0.05 after 5 min exposure, Fig. S2A, B), although not after 20 min exposure (Fig. 1E-F; Fig. S2A).

**Figure 1.**
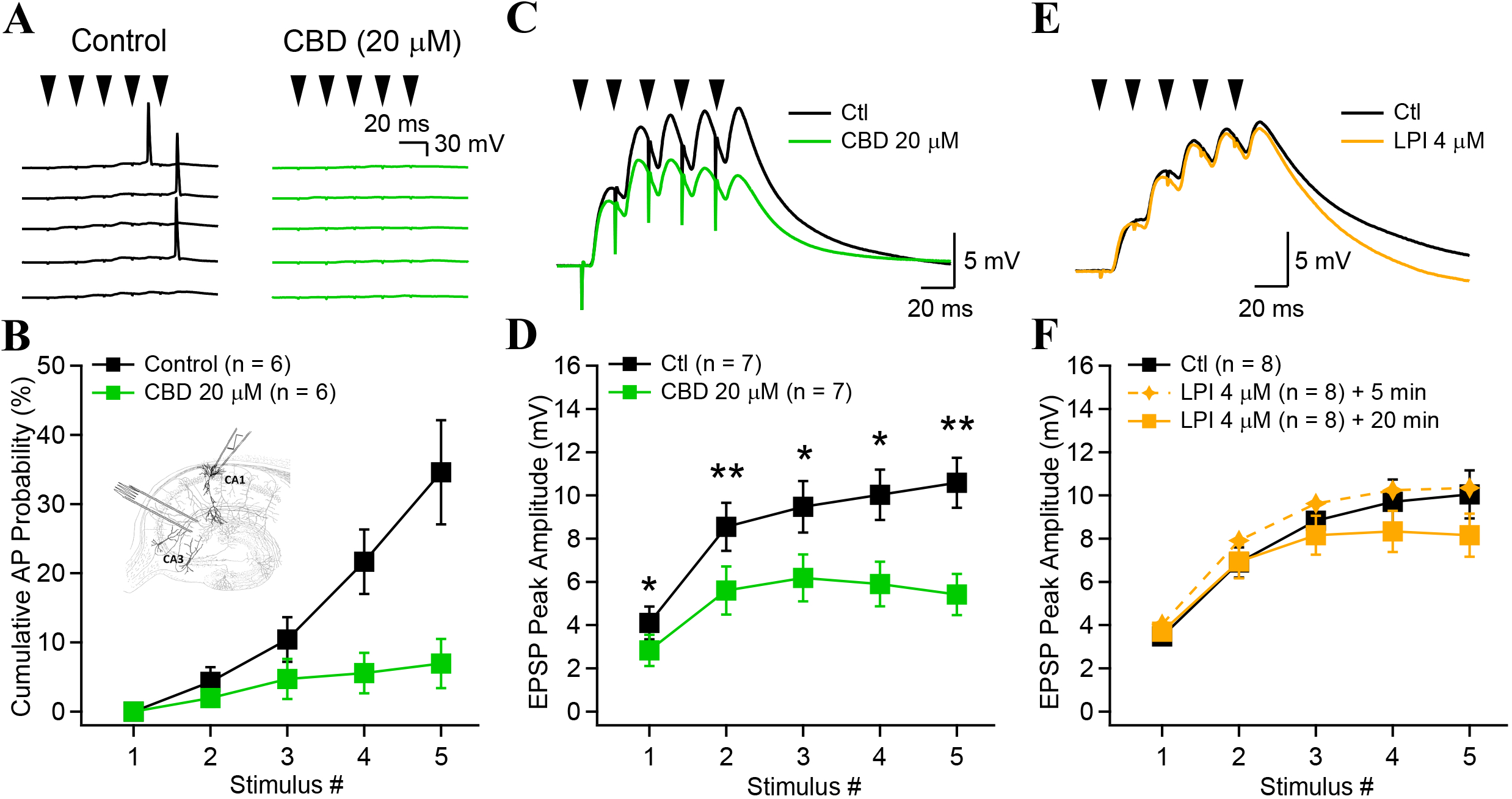
Cannabidiol reduces the propagation of high-frequency activity in the CA1 hippocampus. **A**, Synaptically-evoked action potentials in CA1-PYRs in control conditions (black). CBD decreased AP generation in CA1 pyramidal cells (5 consecutive traces shown for control and CBD). **B**, Cumulative AP probability as a function of stimulus number. Inset, Recording schematic modified from original drawings of Ramon y Cajal showing a recording electrode in a CA1 pyramidal cell and stimulation of Schaffer collaterals with a tungsten electrode. **C**, Compound synaptic potentials recorded in the current-clamp configuration (black trace) evoked by 5 stimulus of Schaffer collaterals delivered at 50 Hz. CBD (20 µM) application for 20 minutes decreased the compound synaptic potential amplitude (green trace). Example traces are the average of 60 consecutive sweeps. **D**, Synaptic potential peak amplitude as a function of stimulus number. CBD significantly decreased the peak synaptic potential for any stimulus number. P-values were corrected for multiple comparison using a post-hoc Holm-Bonferonni correction. **E**, Synaptic potential recorded in control (black) and after application of LPI (orange). **F**, LPI initially increased synaptic potential amplitude but had no long-lasting effect. * p > 0.05; ** p > 0.01. Data points show mean ± SEM.

Second, we tested how CBD affects the intrinsic excitability of CA1-PYRs. CA1-PYRs were challenged with current steps of increasing amplitude to assess AP firing as a function of current injection (F-I curves) before and after CBD application (Fig. 2A-B). CBD potently reduced CA1-PYR firing and capped the maximal firing frequency in response to current injection (Fig. 2A-B). These effects persisted in acute slices from GPR55-KO animals (Fig. 2C-D), suggesting a GPR55-independent effect on CA1-PYRs, consistent with no change in CA1-PYR excitability whether GPR55 signaling varied upward (Rosenberg et al., 2023) or downward (Fig. 2C-D).

**Figure 2.**
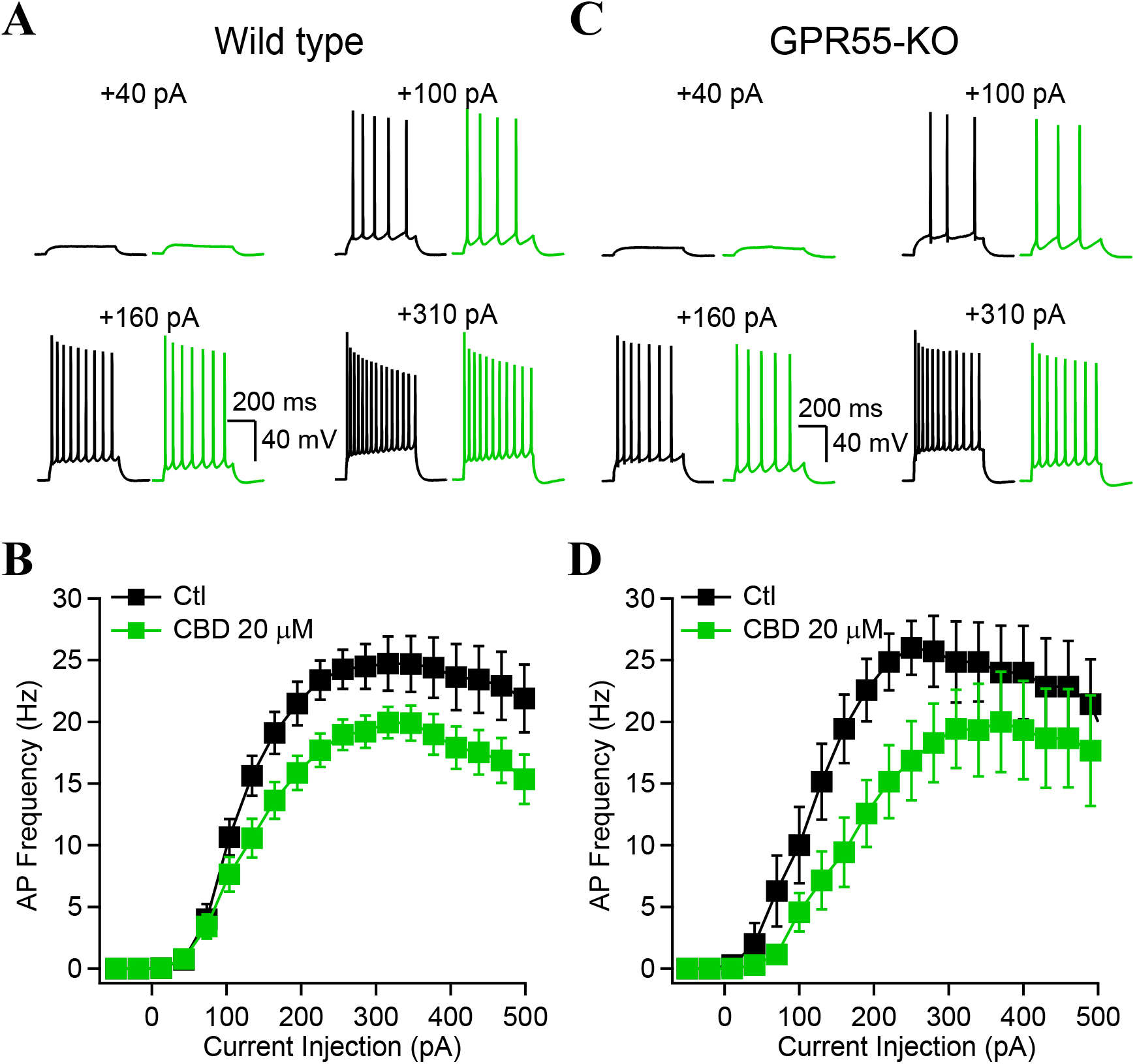
Cannabidiol decreases and caps CA1-PYRs firing independently of GPR55. **A**, Membrane potential and AP firing in response to depolarizing current injected through the recording electrode before (black) and after CBD application (green). **B**, Firing frequency as a function of current injection for n = 16 CA1-PYRs. The FI curves reveal that CBD decreases and caps the firing frequency as a function of current injection. **C**, Similar experiments as in **A** and **B**, but performed in slices from GPR55-KO animals showing similar effect of CBD on the FI curve (**D**). Data points show mean ± SEM for n = 7 CA1-PYRs.

### Cannabidiol requires GPR55 and synaptic inhibition to dampen activity propagation

We next investigated how CBD reduces the propagation of high-frequency activity from hippocampal CA3 to CA1. Although CBD modulation of GPR55 was not implicated in the control of CA1-PYR intrinsic excitability, GPR55 could nonetheless be involved in CBD modulation of synaptic transmission, thereby regulating propagation of high-frequency activity. CBD may act directly on inhibitory interneurons (Kaplan et al., 2017; Khan et al., 2018) which could act as an intermediary onto CA1-PYR to alter spike propagation.

Synaptic potentials recorded in acute slices prepared from GPR55-KO animals displayed similar amplitudes as in wild-type animals (Fig. 3A-B). CBD failed to decrease synaptic potentials evoked in GPR55-KO slices (Fig. 3A-B). We validated the involvement of an antagonistic action of CBD at GPR55 by investigating how a selective GPR55 antagonist (CID-16020046, 10 µM) (Kargl et al., 2013) modulates synaptic potentials. Application of CID significantly decreased synaptic potential amplitudes, similar to CBD’s effects (Fig. 3C-D). Further, CID (20 min) occluded any further effect of CBD, supporting that GPR55 mediates the effects of CBD on synaptic potentials (Fig. 3C-D).

**Figure 3.**
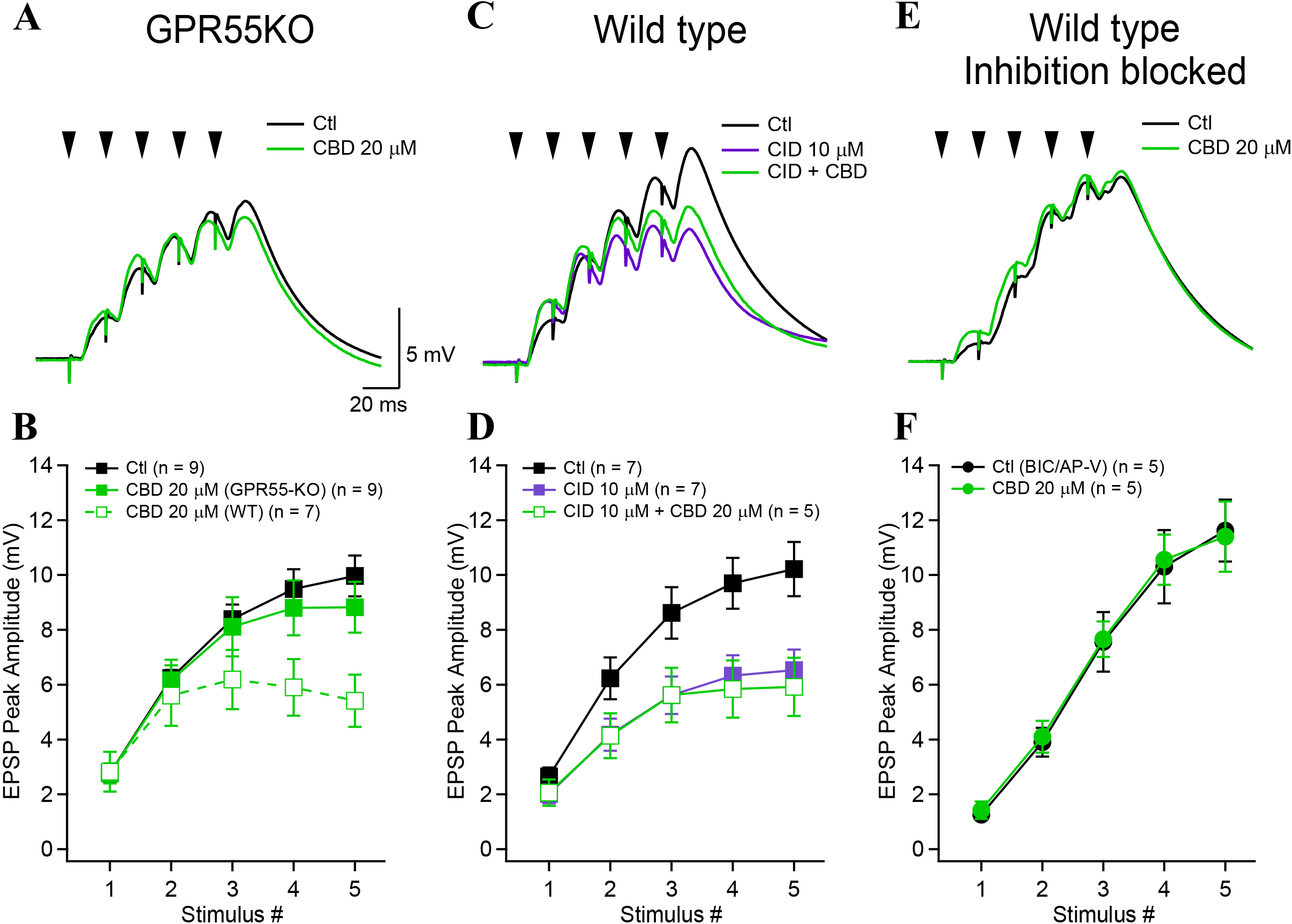
Cannabidiol antagonizes GPR55 and requires synaptic inhibition to decrease the propagation of high-frequency activity. **A**, Synaptic potentials recorded in control (black) and in the presence of CBD (green) for experiments performed in slices obtained from GPR55-KO animals (60 consecutive traces averaged for both conditions). **B**, Synaptic potentials amplitudes as a function of stimulus number. Dotted green trace shows synaptic potential amplitude after CBD in WT animals. **C**, Examples of burst potentials and analysis of their amplitude (**D**) recorded in wild-type animals before (black) and after (magenta) the application of the selective GPR55 inverse agonist CID-16020046 (10 µM) followed by the combination of CID and CBD (green, average of 60 consecutive sweeps shown for all conditions). **E**, Burst potentials recorded in presence of bicuculine (10 µM) and AP-V (50 µM) to block GABA_A_ receptors and NMDAR, respectively. Experiments were performed in WT animals. **F**, Group data from 5 neurons shows that CBD had no effect on synaptic potentials when GABA_A_ receptors were blocked, indicating the CBD requires intact synaptic inhibition to decrease burst potential amplitude. Data points show mean ± SEM.

A decrease in synaptic potential amplitudes could result from a reduced excitatory synaptic transmission or an elevated inhibition synaptic transmission. We tested the idea that CBD modulates inhibitory synaptic transmission. Synaptic potentials were evoked in control condition, followed by bath application of bicuculine (Bic, 10 µM). The NMDAR antagonist AP-V was co-applied to reduce recurrent activity generation. Co-application of Bic and AP-V significantly increased the amplitude of synaptic potentials, evoking action potentials in CA1-PYRs. This confirms that inhibitory transmission influences net synaptic potentials, with maximal inhibition at the end of the train (Fig. S3A-C). Stimulation intensity was adjusted to reduce AP firing, retaining high amplitude synaptic potentials similar to control condition (Fig. 3E-F). CBD had no effect on synaptic potentials recorded in CA1-PYRs with blockade of inhibition (Fig. 3E-F). These results indicate that CBD acts on synaptic inhibition in a GPR55– and inhibition-dependent manner to reduce propagation of high-frequency activity.

### Cannabidiol enhances recruitment of PV-INs but decreases recruitment of SST-Ins

Finding a GPR55-mediated modulation of inhibition raises the question of whether CBD affects recruitment of inhibitory neurons (INs). Parvalbumin-expressing and somatostatin-expressing inhibitory interneurons (PV-INs and SST-INs respectively), are two major IN types accounting for most INs in the CA1 deeper layers that could help mediate CBD’s effect.

Immunohistochemistry revealed that GPR55 was expressed by most PV-INs but only half of SST-INs (Fig. 4A, B), with different colocalization across CA1 strata (Fig. 4B). PV-INs were recruited transiently at the onset of Schaffer collateral stimulation, while SST-INs were recruited later during trains (Fig. 4C-D; Pouille and Scanziani, 2004; Chamberland, Grant, 2024). CBD markedly increased the firing probability of PV-INs while decreasing the firing probability of SST-INs (Fig. 4C-D). CBD largely spared the contrasting dynamics of PV– and SST-INs recruitment: PV-INs remained preferentially active at the train onset, while SST-INs were recruited later and persistently during trains. Following CBD treatment, PV-INs effects dominated throughout the stimulation trains. Recordings of the underlying EPSPs during burst stimulation patterns revealed that CBD had no effect on EPSPs recorded in PV-INs, but strongly decreased EPSPs recorded in SST-INs (Fig. 4E-F). Diminished excitatory drive could explain the decreased firing of SST-INs and is consistent with a general dampening of CA1-PYRs firing and less disynaptic recruitment of SST-INs in the CA3-CA1 circuit. Alternatively, the lack of change in EPSPs in PV-INs does not account for their enhanced recruitment after CBD is given.

**Figure 4.**
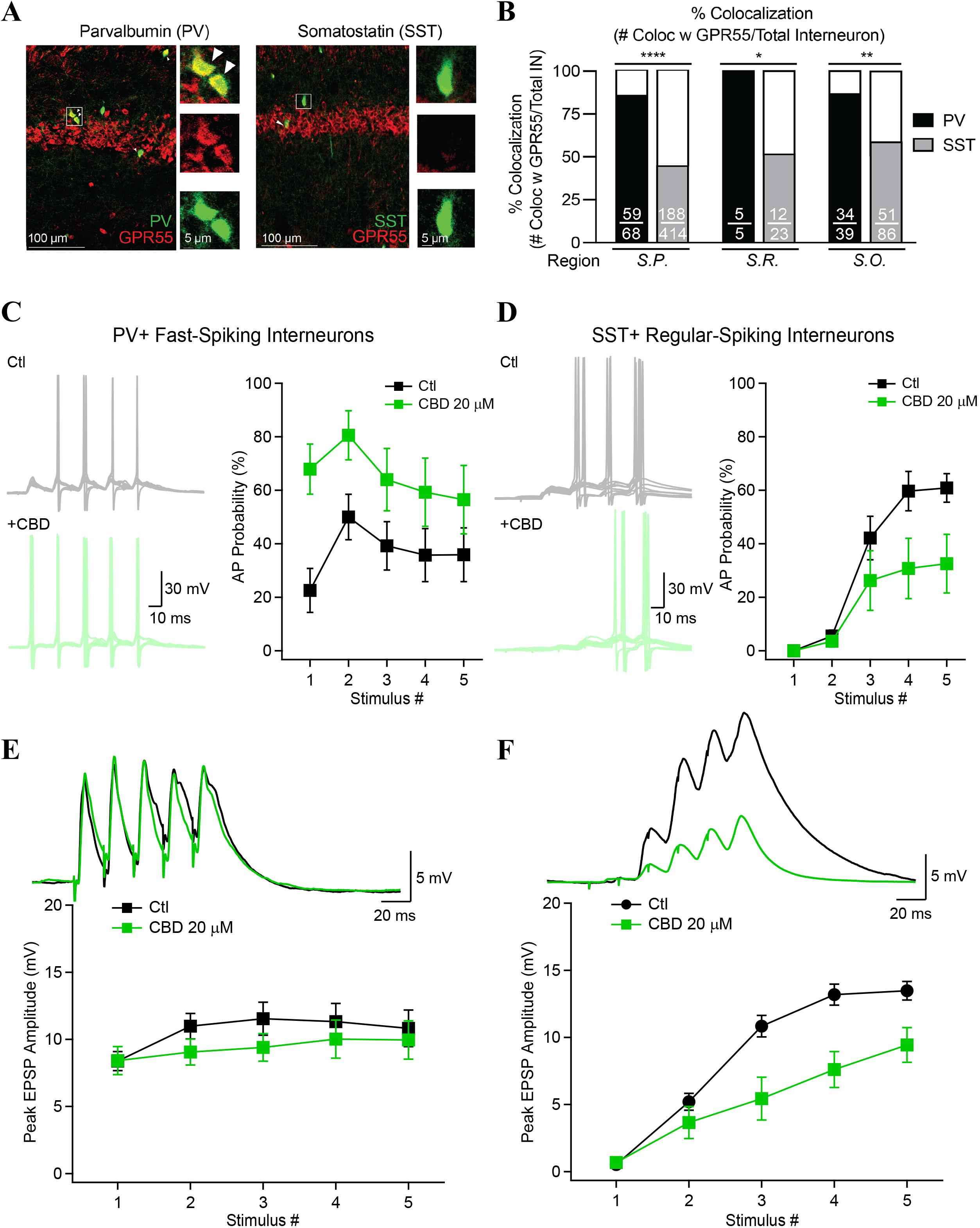
Cannabidiol increases the recruitment of PV-INs but decreases the recruitment of SST-INs. **A**, Confocal images in the CA1 hippocampus showing colocalization between GPR55 (red) and PV (green, left) or SST (green, right) using sections from PV– and SST-Ai9 mice, respectively. Insets denote representative cells as labeled by white box. Arrowheads represent colocalized cells. **B**, GPR55 colocalizes more with PV-INs than SST-INs (n = 2-3 slices each from n = 3 age– and gender-matched mice from each genotype (n=7 total slices). Fractions in bar graphs represent the number of GPR55 co-expressing cells divided by the number of INs. Results analyzed via Chi Squared comparisons between genotypes: S.P. p < 0.0001; S.R. p = 0.0472; and S.O. p = 0.002. **C**, Current-clamp recordings from fast-spiking PV-INs and regular-spiking SST-INs (identified *in situ* by the expression of TdTomato) (**D**) during 5X50 Hz stimulation of Schaffer collaterals. Action potential likelihood as a function of stimulus number is shown. **E**, **F**, Synaptic potentials recorded in PV– and SST-INs in control (black) and after the application of CBD.

### Cannabidiol differentially modulates the intrinsic excitability of PV– and SST-INs

The above results raise the possibility that CBD directly modulates the intrinsic excitability of PV-INs. Neuromodulators such as oxytocin and cholecystokinin, working through G protein coupled receptors, can enhance PV-INs firing in the CA1 hippocampus (Foldy et al., 2007; Owen et al., 2013). This poses the question of whether CBD affects the intrinsic excitability of PV– and SST-INs.

PV– and SST-INs were depolarized with current steps of increasing amplitude to measure F-I curves before and after CBD (Fig. 5). Application of CBD caused a leftward shift in the FI curve of PV-INs (Fig. 5 A-B), but a rightward shift in the F-I curve of SST-INs (Fig. 5C-D). CBD significantly decreased the rheobase measured in PV-INs and shifted the AP threshold towards more negative potential. This contrasted with SST-INs, for which CBD elevated the rheobase, an effect somewhat offset by a slightly more negative AP threshold. The V_M_ and the spontaneous firing rate of PV– and SST-INs were not altered by CBD application (Fig. S4). In this respect, CBD effects on PV-INs differ from those of peptide neuromodulators such as oxytocin (Owen et al., 2013) or CCK (Foldy et al., 2007). PV– and SST-IN activities are time-locked to train stimulation even as their recruitment is differentially altered. Overall, CBD favored the recruitment of PV-INs, by augmenting their intrinsic excitability, but also opposed the recruitment of SST-INs, by lowering their received excitatory synaptic drive (Fig. 4F) (Adaikkan et al., 2024) and by dampening their intrinsic excitability.

**Figure 5.**
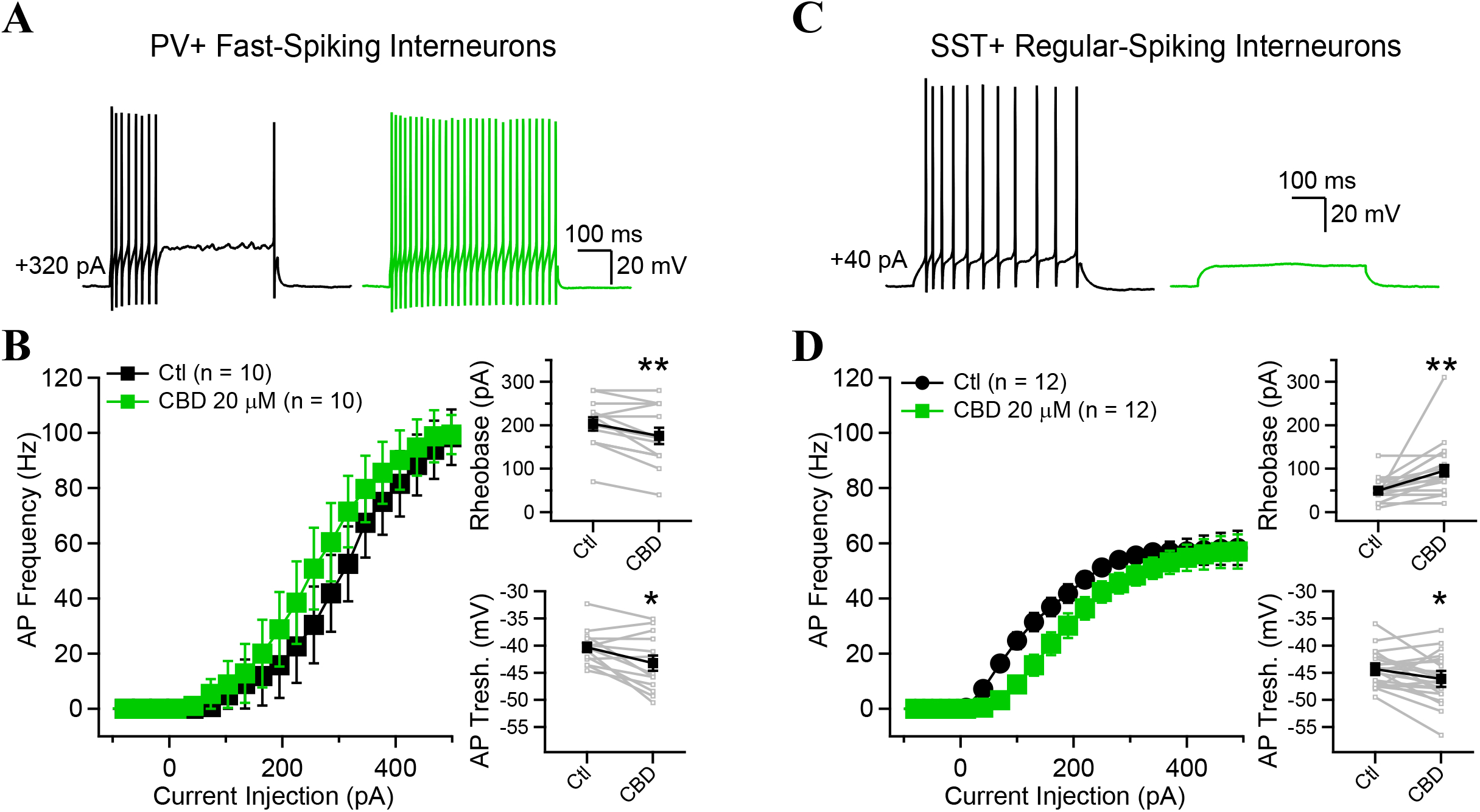
Cannabidiol increases PV-INs but decreases SST-INs firing in response to depolarizing current. **A**, **C**, Examples of AP firing in response to current injection before (black) and after the perfusion of CBD (green). **B**, **D**, FI curves showing the AP probability as a function of current injection in PV– and SST-INs. Top insets show a significant decrease in rheobase for PV-INs but a significant increase for SST-INs. Bottom insets show a more hyperpolarized AP threshold for both types of INs, with a more pronounced decrease observed in PV-INs. * p > 0.05; ** p > 0.01. Data points show mean ± SEM.

### CBD increases feedforward but decreases feedback synaptic inhibition

Since CBD differentially modulates PV– and SST-INs and temporally shifts their recruitment, we asked if CBD alters feedforward and feedback inhibition in the CA1 circuit (schematized in Fig. 6A). Whole-cell voltage-clamp recordings were performed in CA1 PYRs held at depolarized potentials (–40 mV or 0 mV), and the stimulation intensity was adjusted to evoke both polysynaptic feedforward (FF) and feedback (FB) IPSCs (Fig. 6A-B). FF and FB IPSCs were distinguished by their different latencies (horizontal bars, Fig. 6B-C); the difference in latency (on average, 6.3 ms) was consistent with the temporal delay recruiting PV– and SST-INs firing after SC stimulation. The rise time of the FF IPSC (∼1 ms, Fig. 6D) was briefer than the rise time of the FB IPSC (∼2 ms); this is as expected given that the FF IPSC is generated by perisomatic GABA_A_Rs, while the FB IPSC is generated at dendritic locations, electrotonically far from the soma. The multi-synaptic nature of IPSCs was confirmed by applying NBQX and APV, which abolished the inhibitory currents (Fig. 6F).

**Figure 6.**
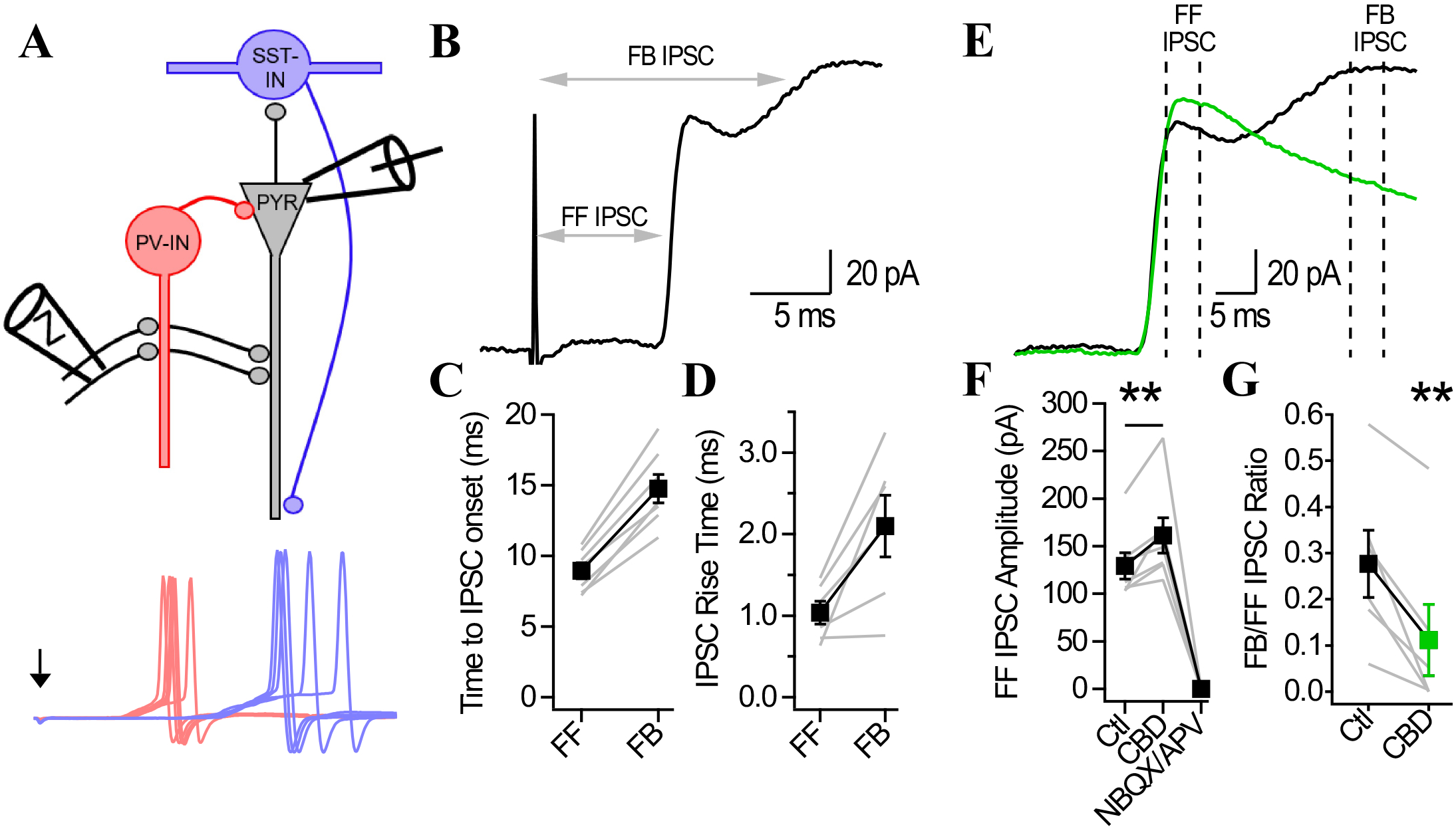
Countermodulation of feedforward and feedback inhibition by cannabidiol. **A**, Schematic of the CA1 microcircuit under investigation. PV-INs are shown in red, while SST-INs are in blue. Bottom illustrates the temporal delay in AP generation observed in PV– and SST-INs. **B**, IPSCs recorded in voltage-clamped CA1 pyramidal neurons held at 0 mV. Note the appearance of a second peak riding on top of the first IPSC. The second IPSC occurred with a time delay corresponding to the recruitment of disynaptic interneurons. The second IPSC had a slower rise time than the first IPSC, hinting at its dendritic origins. **C**, Time from stimulus artifact onset to feedforward and feedback IPSC peaks. **D**, IPSC rise time for the feedforward and feedback IPSCs. **E**, IPSCs recorded at 0 mV in CA1-PYRs before (black) and following CBD application (green). **F**, The amplitude of the FF IPSC was augmented by CBD and both IPSCs were fully abolished by the application of APV/NBQX. **G**, The amplitude of the second IPSC (measured from the trough to the peak) was significantly decreased following CBD application and the FB/FF IPSC ratio was therefore decreased by CBD.

CBD increased the amplitude of the feedforward IPSC (Fig. 6E, F; p < 0.01) but decreased the amplitude of the FB IPSC (Fig. 6E); CBD decreased the ratio of feedback to feedforward IPSC amplitudes in every cell (Fig. 6G). In contrast, in CA1-PYR recordings in slices from PV-Ai32 animals expressing ChR2 in PV interneurons, CBD had no effect on optogenetically-evoked IPSCs (Fig. S5) (Rosenberg et al., 2023). Similarly, CBD had no impact on the frequency or amplitude of spontaneous IPSCs recorded in CA1-PYRs (Fig. S6). These results indicate that CBD strongly biases the local inhibitory network to enhance feedforward perisomatic inhibition in CA1-PYRs, particularly when synaptic afferents recruit interneuron firing, which strongly increases stimulus-locked feedforward inhibition.

### Na_V_1.1 channel enhancers mimic and occlude CBD action on EPSP trains

Dravet Syndrome (DS) usually results from loss-of-function mutations in the *SCN1A* gene encoding the voltage-gated sodium channel Na_V_1.1, a major contributor to excitability in PV and SST INs (Rubinstein et al., 2015). CBD increases PV-INs excitability in different pre-clinical models, including a DS mouse model (Kaplan et al., 2017; Khan et al., 2018). To test if an increase in Na_V_1.1 function, and secondarily PV-INs activity, is a key aspect of CBD action, we combined pharmacological and optogenetic manipulations (Fig. 7 and 8).

**Figure 7.**
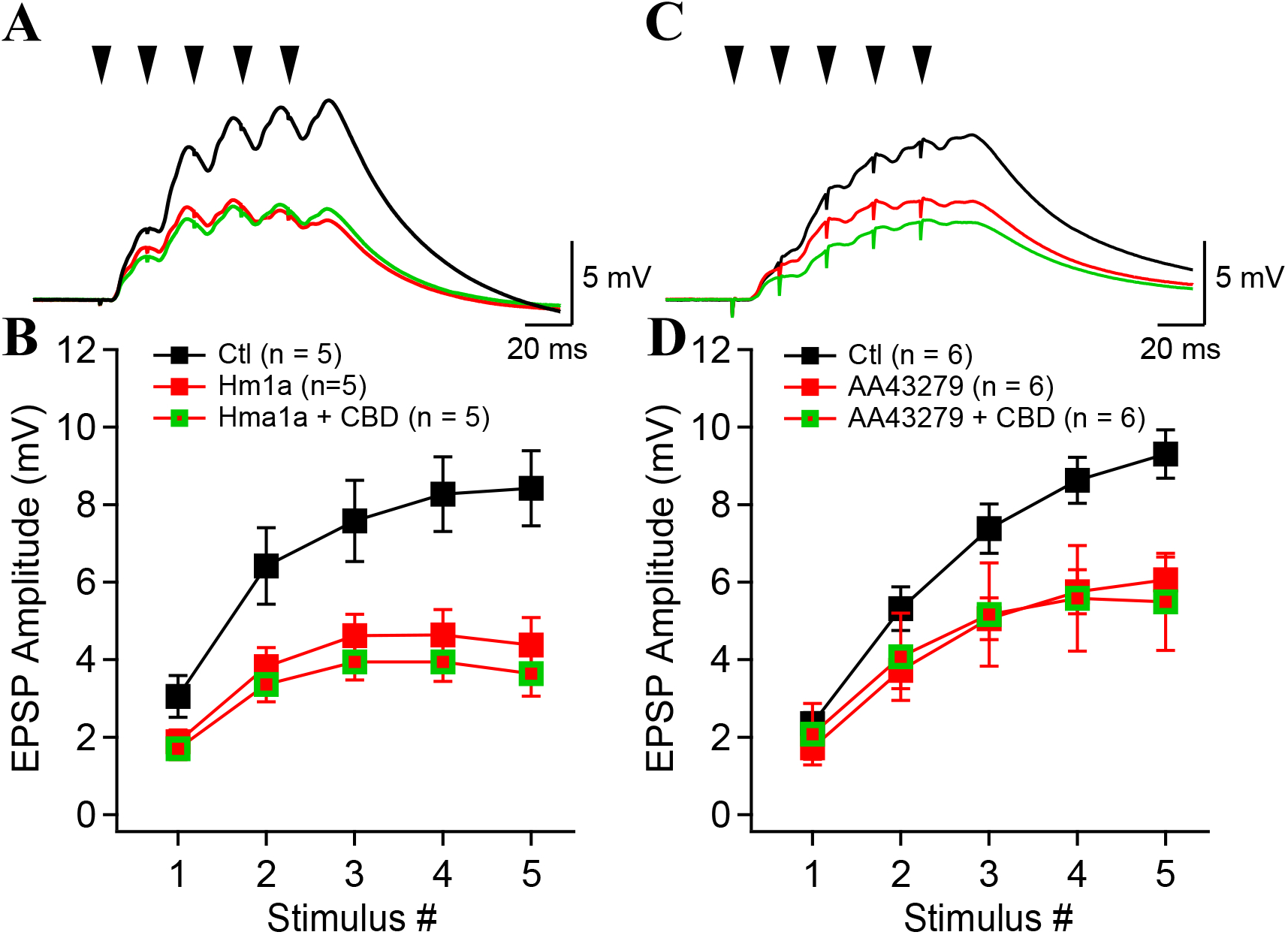
Two Na_V_1.1 enhancer mimic the circuit effects of cannabidiol. **A**, **C**, Examples of synaptic potentials evoked by Schaffer collateral stimulation in control condition (black), in presence of Hm1a (red, A) or AA43279 (red, C) and following the addition of CBD (green). **B**, **D**, Synaptic potential amplitude as a function of stimulus number. Colors are as in A and C.

**Figure 8.**
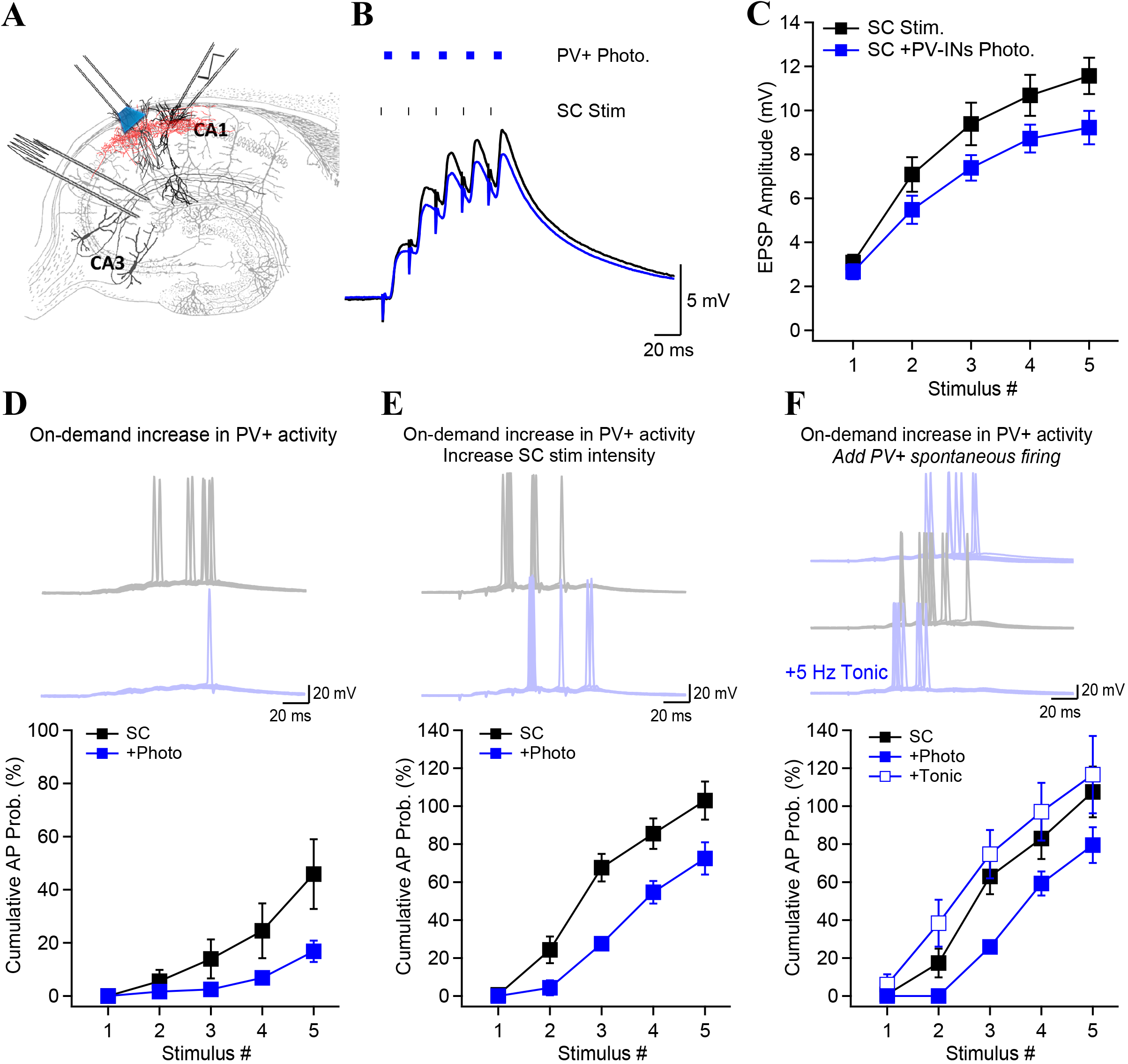
*On-demand* PV+ interneurons firing reduces the propagation of activity. **A**, Experimental design and recording scheme for experiments performed in acute slices from PV-Ai32 mice. **B**, Synaptic potentials recorded in control condition (black) and together simultaneously with blue light photoactivation of PV-INs. **C**, Amplitude of synaptic potentials as a function of stimulus number for control (black) and PV-INs photostimulation (blue). **D**, The stimulation intensity was adjusted to generate APs in control condition. Photoactivation of PV-INs decreased AP probability. Examples show 10 consecutive trials, with control condition in gray and with photostimulation in blue. **E**, The AP probability was doubled compared to D by increasing the electrical stimulation intensity. Photoactivation of PV-INs shifted the generation of APs to later stimuli, and caused a significant decrease in the cumulative AP probability for all stimulus number. **F**, Spontaneous PV-IN firing was mimicked with blue light continuously delivered at 5 Hz. Unlike the on-demand model, spontaneous PV-IN firing caused a leftward shift in the cumulative AP probability, indicative of an increased AP probability earlier in the train. Examples shown in **D**, **E**, **F** are from 10 consecutive trials. Data points show mean ± SEM.

First, we enhanced Na_V_1.1 using Hm1a or AA43279, two molecules that potentiate Na _V_1.1 currents in cell lines and promote PV-IN firing (Frederiksen et al., 2017; Richards et al., 2018) (Fig. 7). Application of either Hm1a or AA43279 significantly decreased the synaptic potential amplitude in CA1-PYR (Fig. 7A-D), similarly to CBD (Fig. 1C-D). Further, Hm1a and AA43279 occluded any further effect of CBD on synaptic potentials (Fig. 7A-D), suggesting converging mechanisms of action.

### Specific, stimulus-locked activation of PV-INs recapitulates the effects of CBD

Second, we used optogenetics to test how directly augmenting FFI impacts the propagation of activity from CA3 to CA1 (Fig. 8). Blue light illumination of slices from PV-Ai32 animals expressing ChR2 in PV interneurons enabled stimulation of PV-INs with cellular specificity and temporal precision (Fig. 8A). Compound postsynaptic potentials evoked by Schaffer collaterals stimulation were significantly decreased when optogenetic stimulation of PV-INs was co-applied with electrical stimulation in a time-locked fashion (Fig. 8A-C). The reduction in net EPSP was fully reversible (Fig. S7). The firing likelihood was decreased by time-locked optogenetic activation of PV-INs (Fig. 8D), recapitulating CBD’s effect (Fig. 1A-B). The decrease in AP likelihood was maintained even after the electrical stimulation intensity of Schaffer collateral afferents was increased (Fig. 8E). In contrast, photostimulation of SST-INs exerted no such effect on synaptic potentials and AP probability (Fig. S8). Therefore, pharmacological enhancement (Fig. 7A-D) or stimulus-locked activation of PV-INs (Fig. 8) are each sufficient to decrease the propagation of high-frequency activity from CA3 to CA1, even under high levels of activity.

We predicted that enhanced excitability of PV-INs, likely mediated by increasing Na_V_1.1 activity, would increase perisomatic inhibition when it is most needed, just after feedforward excitation. To explore the concept of such increased perisomatic inhibition “on-demand”, we compared the effects of precisely timed optogenetic stimulation with the outcome of driving PV-INs to fire continuously, as if under the influence of neuromodulators like oxytocin or CCK (Foldy et al., 2007; Owen et al., 2013). Such continual firing of PV-INs was mimicked by tonically photostimulating PV-INs afferents at 5 Hz (Fig. 8F). Under this condition, optogenetic stimulation of PV-INs failed to dampen the AP likelihood but produced instead a leftward shift in the stimulus number-dependent probability of firing (Fig. 8F). This was consistent with Owen et al’s (2013) findings that continuous recruitment of PV-INs by oxytocin or CCK causes fatigue at their inhibitory outputs on PYR neurons, favoring the propagation of activity in hippocampal CA1 rather than dampening it (Owen et al., 2013). This reinforces that feedforward perisomatic inhibition is most effective upon precisely timed delivery in concert with excitation. CBD accomplishes this by enhancing the on-demand recruitment of PV-IN firing while preserving its temporal precision.

## Discussion

We uncovered a novel, multipronged mechanism by which CBD alters propagation of high-frequency activity in the hippocampus, simultaneously (1) decreasing principal neuron activity, (2) enhancing FF inhibition and (3) reducing delayed FB inhibition. Our dissection of CBD’s actions revealed that it changed the excitability of constituent neurons in different directions, reducing the excitability of CA1-PYRs, increasing the excitability of PV-INs, but concomitantly dampening the excitability of SST-INs. This is a more nuanced outcome than simply decreasing excitation and increasing inhibition, the prevailing conception of an ideal anti-seizure medication (ASM). Our experiments illuminate how CBD’s disparate effects on individual neuronal types influence its efficacy. Better defining microcircuit impacts could improve how CBD and other ASMs are used to treat diverse epilepsies. ASM resistance is a worldwide problem (World Health Organization, 2019). Voltage– and ligand-gated ion channels are the leading ASM mechanisms of action but are not confined to excitatory neurons or inhibitory neurons. This holds for common substrates of ASM action including sodium, calcium, and potassium channels as well as receptors for GABA and glutamate (Rogawski, 2013). The beneficial action of an ASM, mediated by quieting glutamatergic principal neurons, can be eroded by its deleterious proconvulsant effect on GABAergic interneurons. This was rigorously demonstrated for a K_V_7 potassium channel activator, retigabine (Jing et al., 2022), showing how yoked actions on excitatory and inhibitory neurons can partially cancel each other.

While epilepsy may originate from malfunction of individual channel types, it invariably manifests at the neuronal circuit level. Considerations of canonical brain circuits raise challenges for seizure control and offer a way forward. Paz and Huguenard (2015) have summarized basic microcircuit motifs broadly expressed in brain networks and pointed out how their selective manipulation could quell ictal propagation. For example, microcircuits for feed-forward inhibition exist in hippocampus, neocortex and thalamus could provide strategic chokepoints for an ASM to prevent of ictal propagation. In contrast, the impact of upregulation of microcircuits for feed-back inhibition was less clear, with potential favorable, mixed, or unfavorable effects (Paz and Huguenard, 2015). Expectations about microcircuit effects can be evaluated preclinically with optogenetic and pharmacogenetic manipulations to clarify the effects of IN types and microcircuits (Shiri et al., 2017; Bohannon and Hablitz, 2018; Wang et al., 2020; Lado, Xu and Hablitz, 2022).

### Effects of CBD on specific hippocampal neurons and subcircuits

Following the initial demonstration that CBD exerts antiepileptiform and antiseizure properties not mediated by conventional cannabinoid receptors (Jones et al., 2010), key studies focused attention on the role of synaptic inhibition and increases in the excitability of fast-spiking inhibitory neurons (Kaplan et al., 2017; Khan et al., 2018). We confirmed the experiments of Kaplan *et al*. and Khan *et al*. on CBD-enhancement of fast-spiking neurons’ firing (Fig. 5A, B) and demonstrated that CBD has opposing effects on CA1-PYRs and SST-INs. Prior CBD studies mainly relied on recordings of spontaneous IPSCs in PYR neurons, which could not differentiate between feedforward and feedback inhibition. We separated these targets and found that CBD strengthens FF inhibition and dampens FB inhibition (Fig. 6). This pattern of changes in the CA1 circuit is more complex than envisioned for an ASM that simply renormalizes E-I imbalance. We now discuss possible mechanisms for individual cell types and inhibitory motifs and implications for the circuit as whole.

#### PV-INs and feedforward inhibition

Our experiments narrow down the signaling mechanism for CBD-induced enhancement of feedforward inhibition. Effects of CBD were mimicked by a GRP55 agonist CID (Fig. 3C, D) and eliminated by GPR55 deletion (Fig. 3A, B), strongly implicating GPR55 signaling. Effects of CBD were not explained by an enhancement of synaptic inputs onto PV-INs (Fig. 4E) or on their synaptic output strength (Fig. S5). These findings strongly implicate direct effects of CBD on the intermediary PV-IN excitability (Fig. 5 AB). The biophysical basis of the augmented excitability remains unclear, however could theoretically involve effects on either Na or K channel targets. Supporting a role for Na channels, pharmacological activation of Na_v_1.1 (Fig. 7), was sufficient to mimic and occlude effects of CBD on feedforward inhibition. These channels are highly enriched in PV-INs (Ogiwara et al., 2007). One hypothesis suggests that CBD could block basal activity of the GPR55 agonist LPI (Kapur et al., 2009), thereby relieving PKC activation, PKC-mediated α_1_ subunit phosphorylation and basal Na^+^ channel suppression (Cantrell and Catterall, 2001). Blockade of GPR55 by CBD could disinhibit spike properties. This hypothesis invokes basal GPR55 signaling, plausible when CBD action is studied under seizure-inducing conditions (Kaplan et al., 2017; Khan et al., 2018) or with 50 Hz spike trains (this study), that drive presynaptic LPI release (Sylantyev et al., 2013). These findings predict that effects of CBD on PV-INs excitability would be more pronounced with activity dependent LPI release (Sylantyev et al., 2013; Rosenberg et al., 2023). While K_v_7 has been suggested as a potential target for CBD (Lauckner et al., 2008), prior reports predict that CBD would block enhanced K_V_7-based M current through GPR55, producing a *decrease* in PV firing which opposes the *increase* in PV-IN firing shown here.

Regardless of biophysical mechanism, elevated recruitment of PV-INs in a stimulus-locked manner potently curbs propagation of epileptiform activity. Our experiments reinforce how PV-INs powerfully modulate neuronal networks (Hu, Gan and Jonas, 2014), holding the line between physiological hippocampal oscillations (e.g., sharp-wave ripples) and pathological epileptiform activity (Karlocai et al., 2014). Our *in vitro* optogenetic experiments complement the demonstration that optogenetic activation of PV-INs can stop ongoing electrographic and behavioral seizures (Krook-Magnuson et al., 2013). CBD captures the on-demand aspect of optogenetic stimulation in elevating the responsiveness of PV-INs. The feedforward inhibitory network broadly disperses the dampening of spike propagation, avoiding the pro-epileptic consequences of activation of multiple PV-INs in a restricted space, which can drive rebound spiking in PYR neurons and create a focal seizure point (Sessolo et al., 2015). CBD’s action on PV-IN excitability also stops short of driving PV-INs to fire out-of-synch with the rest of the network. This would occur during spontaneous firing driven by G protein-coupled neuromodulation by oxytocin (Owen et al., 2013) and cholecystokinin (Foldy et al., 2007), which fatigues GABAergic transmission. Simulating out-of-synch firing, we found weakening of CA1 chokepoint function (Fig. 8F), unlike CBD’s beneficial effect.

#### Pyramidal neurons and monosynaptically driven spike throughput

Our recordings demonstrated another CBD action, independent of GPR55: the attenuation of PYR neuron excitability (see also Khan et al., 2018), which was unaffected by GPR55 deletion (Fig. 2). Others have documented CBD’s direct interactions with molecules supporting intrinsic conductances, including direct effects on sodium channels (Ghovanloo et al., 2018; Sait et al., 2020; Zhang and Bean, 2021; Ghovanloo and Ruben, 2022). We proposed a pathophysiological scenario in which circuit overactivity promotes GPR55 receptor expression in a positive feedback loop (Rosenberg et al., 2023; Tsien and Rosenberg, 2025). Here, dampening of PYR cell firing (Fig. 2) is synergistic with GPR55-dependent shifts in E:I ratio (Fig. 3). These two CBD effects are reinforcing, favoring CA1’s capability as a chokepoint for hyperactivity (Paz and Huguenard, 2015). Combining GPR55-dependent and GPR55-independent actions may not be attained by a synthetic GPR55 antagonist like CID16020046.

#### Somatostatin-INs and feedback inhibition

Our observations on neuronal firing and circuit properties aligned as CBD rendered SST-INs less prone to fire, weakening feedback inhibition. Some attenuation of feedback inhibition is expected from lessened PYR neuron firing. The contrast with PV-INs may be surprising given commonalities between SST– and PV-INs excitabilities. For example, the impairment produced by deleting Na_V_1.1 in SST-INs is similar to that seen in PV-INs (Rubinstein et al., 2015). However, the disparities we observed in PV– and SST-INs given CBD are similar to Khan, Ali et al. (2018), who first reported that PV+ and CCK+ basket cells responded differently to CBD (Khan et al., 2018). This underscores the need for caution in viewing all inhibition as a bloc or generalizing effects on different interneuron types (Liebergall and Goldberg, 2024) and circuit motifs (Paz and Huguenard, 2015). The contrasting effects with CBD raise questions about mechanism and therapeutic consequences.

We lack an understanding of why CBD renders PV-INs more excitable and SST-INs less so. The increased PV-IN excitability mediated by GPR55 may reflect higher GPR55 expression in PV-INs than in SST-INs (Fig. 4). Biophysical mechanisms of excitability also differ: the interneuron types differ in spike amplitude, dynamics of spike generation, and respond differently to introduction of the same genetic variant of Kcnt1, a spike-modulating ion channel (Shore et al., 2024). Consistently, dopamine D2 receptor activation oppositely modulates firing in PV-versus SST-INs within CA1(Tuduri et al., 2025).

Is the action of CBD on SST-INs and feedback inhibition a feature or a bug? Opposing the firing of SST-INs could be beneficial if subpopulations of SST-INs that have disinhibitory effects on PYR firing were dominant. *Sst;;Tac1*-INs overwhelmingly target PV-INs over CA1-PYRs; quieting them would facilitate feedforward inhibition. Likewise, a subpopulation of OLM interneurons identified by the expression of *Sst* and *Chrna2* innervate local INs in *lacunosum moleculare* and disinhibit the integration of excitatory inputs from CA3 (Leao et al., 2012; Artinian and Lacaille, 2018). CBD inhibition of this subclass would also dampen CA1 spike throughput.

Countervailing arguments suggest that the dampening of SST-INs has detrimental effects, masked by CBD’s net positive impact. SST-INs silencing turns up CA1-PYRs firing during bursts (Royer et al., 2012), and silencing of feedback inhibition shifts high-frequency oscillations into a pathological mode (Adaikkan et al., 2024). Dampening of SST-IN firing arises from Na_V_1.1 haploinsufficiency in DS mouse models and is interpreted as pro-seizure (Tai et al., 2014; Rubinstein et al., 2015). Our own optogenetic experiments revealed no significant effect of SST-IN activation on the high-frequency PSPs, unlike PV-IN activation. Counteracting changes in inhibition and disinhibition may cancel each other out.

### Preclinical implications

Defining CBD’s multimodal action on key neurons and neuronal circuits offers insights into how this phytocannabinoid dampens seizures in FDA-approved settings of DS, Lennox-Gastaut Syndrome and Tuberous Sclerosis complex (Devinsky et al., 2017; Devinsky et al., 2018). CBD effects on PYR spike propagation, PV neuron firing, and feedback inhibition involve a beneficial combination of GPR55-independent and GPR55-dependent mechanisms (Rosenberg et al., 2023; Tsien and Rosenberg, 2025), opposing activity-dependent, biochemically-based positive feedback. Two possible therapeutic implications arise. First, the effectiveness of CBD in experimental seizure settings (Rosenberg et al., 2023) would not be matched by pure GPR55 antagonism alone; a synthetic GPR55 antagonist like CID16020046 may not combine GPR55-dependent and GPR55-independent actions, although this needs direct testing. Second, CBD will be of less benefit in pathological settings where firing of PV-INs is already maximal or PV-INs or their axons have died or become non-functional (Sloviter, 1991). Our experiments also suggest that CBD actions on SST-INs are a mixed blessing, attenuating feedback inhibition but possibly blunting SST-based weakening of feedforward inhibition. CBD’s diverse actions reflect its combined effects on neuronal types and inhibitory subcircuits, creating chokepoints to limit the propagation and perpetuation of epileptiform activity.

## Material and Methods

### Acute hippocampal slice preparation

All procedures involving mice were approved by the Institutional Animal Care and Use Committee (IACUC) at New York University Langone Medical Center. C57BL/6 (WT), PV-Ai9, SST-Ai9, PV-Ai32, SST-Ai32 and GPR55-KO mice (P21 – P60) of either sex were used. Homozygous PV-Cre mice (Jackson Labs; Stock No. 008069) (Hippenmeyer et al., 2005) and homozygous SST-Cre mice (Jackson Labs; Stock No. 013044) (Taniguchi et al., 2011) were crossed with homozygous Ai9 mice (Jackson Labs; Stock No. 007909) (Madisen et al., 2010) or homozygous Ai32 mice (Jackson Labs; Stock No. 024109) (Madisen et al., 2012) to generate animals for experiments (Chamberland et al., 2023). GPR55-KO mice (*B6;129S-Gpr55^tm1Lex^/Mmnc*) mice were a gift from Prof. Ken Mackie (Indiana University). Upon receipt at NYU, they were backcrossed to C57BL/6J mice (Jax Strain #000664) to expand the colony. Mice were anesthetized with isoflurane before decapitation. The brain was rapidly extracted and placed in ice-cold (4°C) cutting solution, containing (in mM): sucrose 185, NaHCO_3_ 25, KCl 2.5, NaH_2_PO_4_ 1.25, MgCl_2_ 10, CaCl_2_ 0.5, and glucose 25 (pH = 7.4, 330 mOsm). The brain was hemisected and glued on a specimen disk of a VT1000S vibratome (Leica). Transverse hippocampal slices of 300 µm were cut and transferred to heated (32°C) cutting solution where they were held for 30 minutes before being transferred to heated (32°C) recording ACSF in a submerged chamber. The recording ACSF solution contained (in mM): NaCl 125, NaHCO_3_ 25, KCl 2.5, MgCl_2_ 2, CaCl_2_ 2, and glucose 10 (pH = 7.4, 300 mOsm). The slices were allowed to rest for an extra 30 minutes while the ACSF solution was left to cool down to room temperature before recordings were started. The total recovery time was 1 hour following the slicing procedure and slices were used for up to 6 hours after preparation.

### Electrophysiological recordings

Acute hippocampal slices were transferred to a recording chamber that was continuously perfused (2 ml/min) with oxygenated (95% O_2_/5% CO_2_) recording ACSF. Recordings were performed at room temperature. Slices were held under a nylon mesh. An upright microscope (Olympus, BX61WI) equipped with a 40X water immersion objection (Zeiss, 0.8 NA) was used to visually identify neurons for whole-cell and cell-attached recordings. GABAergic interneurons in the PV-Ai9 and SST-Ai9 transgenic mouse models were identified based on their expression of TdTomato. Deep CA1 pyramidal cells were visually targeted at the border between strata oriens and pyramidale for whole-cell recordings. Borosilicate glass micropipettes (TW150-4, World Precision Instruments) were pulled on a P-97 micropipette puller (Sutter). Micropipettes had a resistance of 3 – 6 MΩ for whole-cell recording and a resistance of 5 – 8 MΩ for cell-attached recordings. Current-clamp and voltage-clamp recordings were performed with a solution containing, in mM: 130 K-gluconate, 10 HEPES, 2 MgCl_2_.6H_2_O, 2 Mg_2_ATP, 0.3 NaGTP, 7 Na_2_-Phosphocreatine, 0.6 EGTA, 5 KCl; pH 7.2 and 295 mOsm. Voltage-clamp recordings were performed with intracellular solution containing: 130 Cs-methanesulfonate, 10 HEPES, 2 MgCl_2_.6H_2_O, 4 Mg_2_ATP, 0.3 NaGTP, 7 diTris-Phosphocreatine, 0.6 EGTA, 5 KCl; pH 7.2 and 295 mOsm. For recordings with high intracellular chloride, the solution contained: 130 Cs-chloride, 10 HEPES, 2 MgCl_2_.6H_2_O, 4 Mg_2_ATP, 0.3 NaGTP, 7 diTris-Phosphocreatine, 0.6 EGTA, 5 KCl; pH 7.2 and 295 mOsm. Series resistance was not compensated. Access and input resistance were monitored throughout recording. Cells in which the access resistance varied by > 20% during the recording were discarded from further analysis. Cell-attached recordings were performed with ACSF in the pipette. Schaffer collaterals were stimulated using a tungsten electrode positioned in *stratum radiatum* of region CA3. The stimulation electrode was connected to a stimulus isolator (A360, World Precision Instruments). The stimulation intensity gradually increased to produce synaptic potentials in CA1 pyramidal neurons. The electrophysiological signal was amplified with an Axopatch 200B and digitized at 10 kHz (Digidata 1322A, Molecular Devices). Data was recorded on a personal computer with the Clampex 8.2 software and analyzed in Microsoft Excel and Igor Pro.

### Optogenetics stimulation

An optical fiber connected to an LED driver was positioned at the border between *strata oriens* and *pyramidale*. A TTL signal was simultaneously sent from the digitizer to the electrical stimulator and a Grass stimulus isolator. The Grass stimulus isolator was connected to the LED driver to provide independent control over the optogenetic stimulation. Blue light (470 nm) was delivered as 5 – 10 ms pulse synchronized with the electrical stimulation. For experiments mimicking the spontaneous activation of PV-INs, blue light was continuously delivered at 5 Hz during the trials, in addition to the synchronized photoactivation of PV-INs with the electrical stimulation.

### Immunohistochemistry and image analysis

Mice were perfused transcardially with a 4% paraformaldehyde solution in PBS. Brains were extracted, dissected, and maintained in 4% PFA overnight. The brains were then protected in sucrose solution (30%) overnight. Brain tissue was embedded in Tissue-TEK O.C.T. compound and frozen before sectioning. Sections (16 μm) were obtained by slicing the brain on a frozen cyrostat (Leica). Sections were collected on microscope slides coated with HistoBond (VWR). Sections were treated with 0.2% Triton X-100 and 5% normal serum for 2-3 hours at room temperature, then incubated at 4°C with 1:500 rabbit anti-GPR55 antibody (Cayman Chemical, #10224). Sections were rinsed and incubated with Alexa Fluor-conjugated secondary antibodies and mounted on microscope slides with ProLong Gold Antifade mounting medium. Images were acquired with 10X, 20X and 63X objectives on a Zeiss LSM 510 confocal microscope. Image analysis was performed using custom analysis scripts in Icy (http://icy.bioimageanalysis.org).

CA1 layers were visually identified. Fluorescence was detected independently for TdTomato (Ai9) and GPR55 immunosignal with automated functions in Icy. The colocalization ratio was calculated as the number of GPR55-labeled cells co-expressing TdTomato (Ai9) divided by the total number of labeled interneurons. Statistical comparisons of colocalized versus non-colocalized cells across genotypes (PV-Ai9, SST-Ai9) were performed using a Chi-squared test.

## Supplementary Figure Legends

**Supplementary Figure 1.**
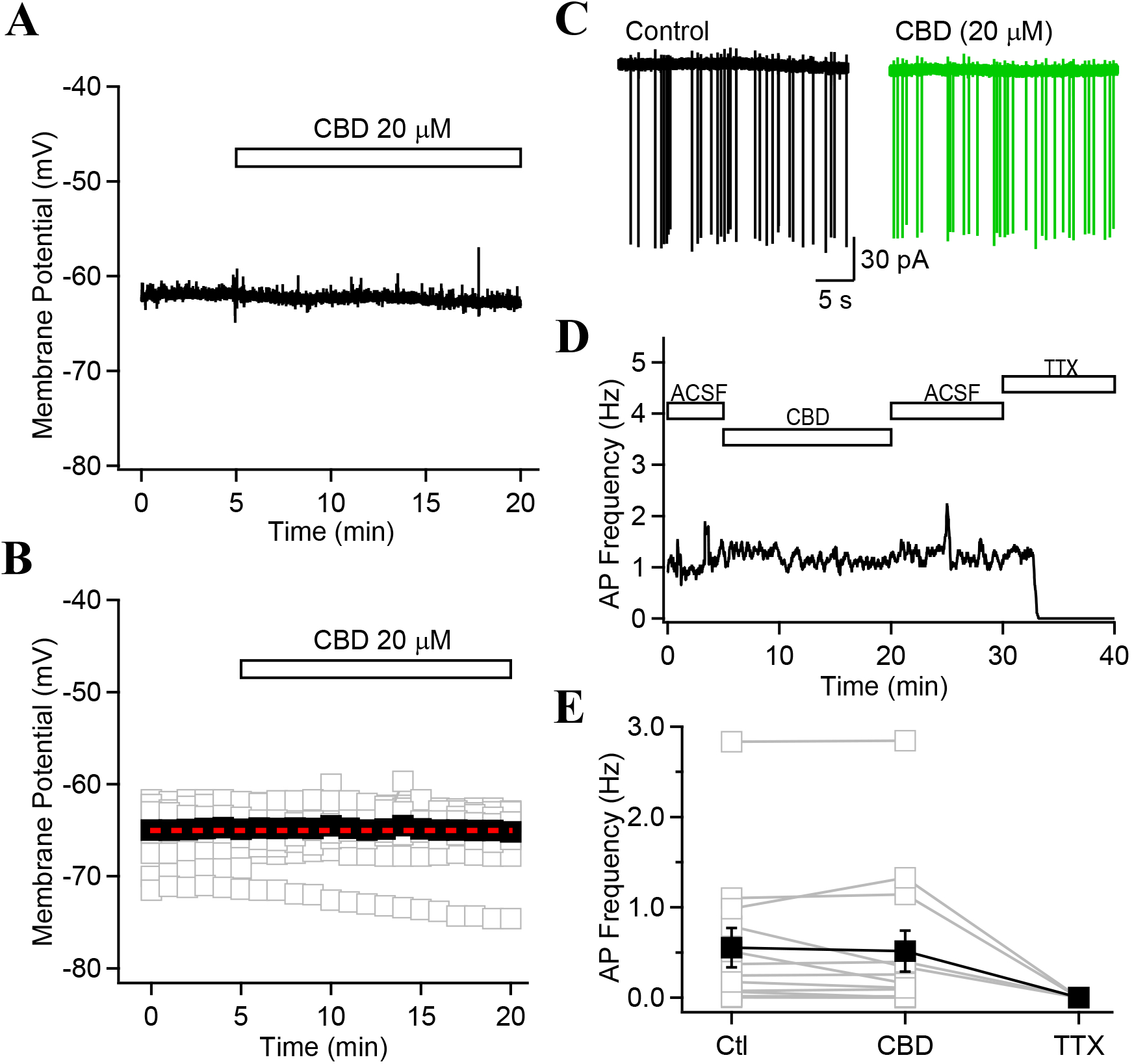
CBD has no effect on the resting membrane potential or the spontaneous firing of CA1-PYRs. **A, B**, Whole-cell recordings showing that CBD has no effect on the resting membrane potential of CA1-PYRs. **C**, Cell-attached recordings of spontaneous AP firing in a CA1-PYR in control (black) and following the continuous application of CBD (green) for 20 minutes. **D**, 40 minutes recording from the examples shown above, in which the spontaneous firing frequency was monitored during CBD application and ACSF wash. No effect of CBD was observed. TTX abolished the inward currents in recorded in cell-attached configuration, confirming that they were APs. **E**, Summary graph showing no change in the spontaneous AP frequency for 13 CA1-PYRs CBD application. 4 neurons were subsequently treated with TTX, which fully blocked all events. Data points show mean ± SEM.

**Supplementary Figure 2.**
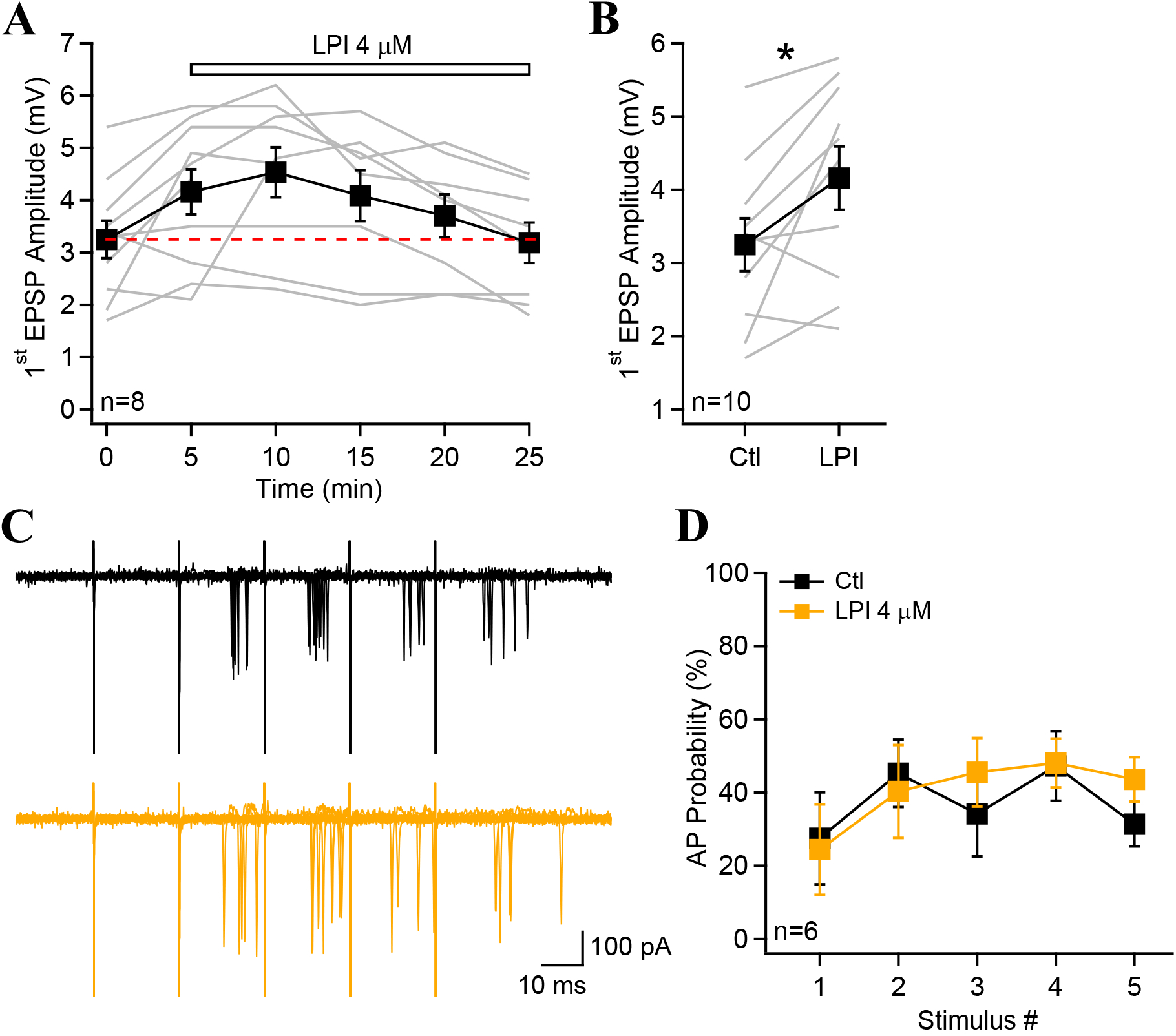
LPI transiently increase synaptic potentials. **A**, Amplitude of the first EPSP as a function of time. Bath application of LPI resulted in a transient increase in the EPSP amplitude that declined over time. **B**, The first EPSP amplitude was significantly increased by 5 minutes LPI application. **C**, Cell-attached recordings from a CA1 pyramidal cell during repetitive Schaffer collaterals stimulation (5X50 Hz) in control (black) and 20 minutes following LPI treatment. **D**, AP probability as a function of stimulus number for group data presented in G. Consistent with whole-cell current-clamp recordings, LPI had no long-lasting effect on the AP probability. Data points show mean ± SEM.

**Supplementary Figure 3.**
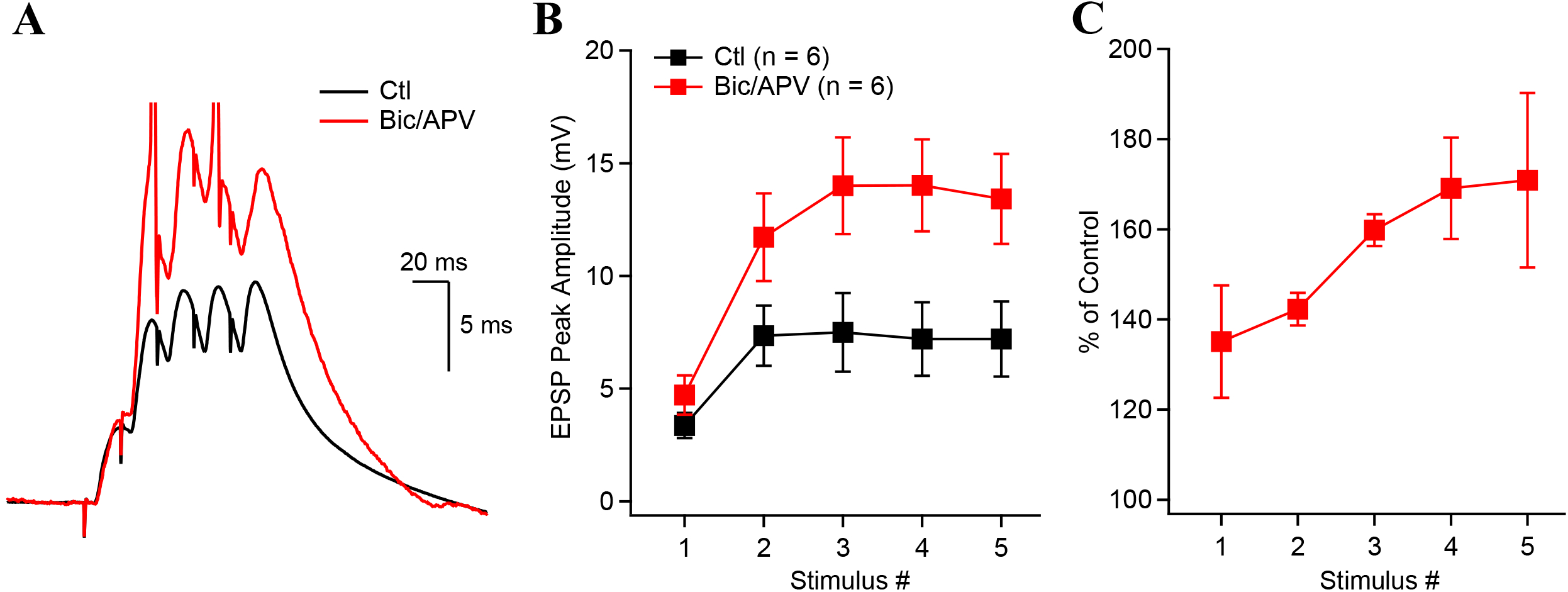
GABA_A_R-mediated inhibition shape synaptic potentials. **A**, Synaptic potentials recorded in control (black) and after the application of bicuculine and AP-V (red). Note truncated action potentials. **B**, Synaptic potential peak amplitude as a function of stimulus number before and after the co-application of bicuculine and AP-V. **C**, Bicuculine had a greater effect on the last synaptic potentials recorded in response to burst stimulation. Data points show mean ± SEM.

**Supplementary Figure 4.**
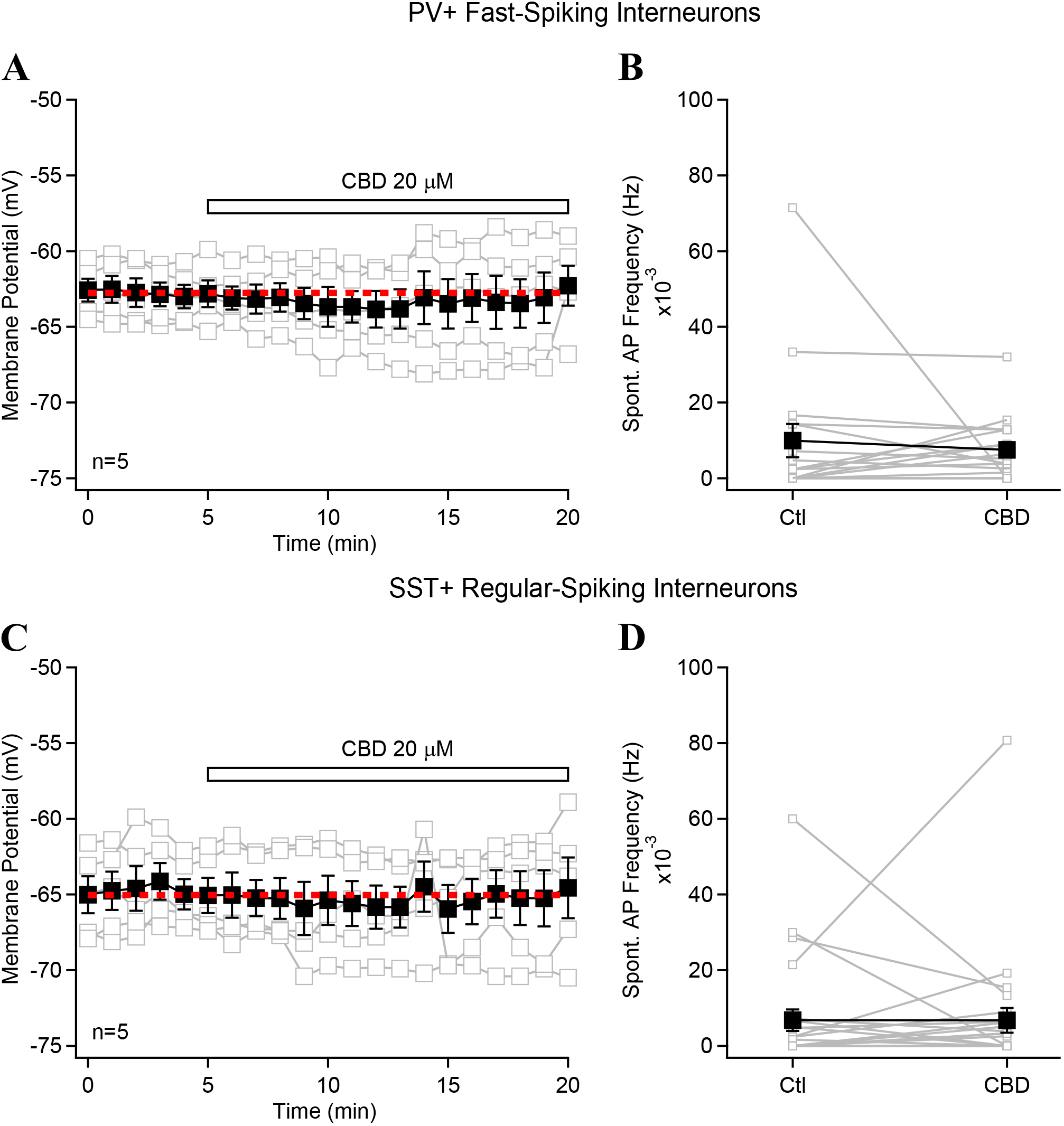
Cannabidiol has no affect the membrane potential or spontaneous firing rate of PV-or SST-INs. **A**, Membrane potential recorded continuously before and during CBD application in PV+ fast-spiking interneurons. Individual neurons are shown in with their average in black and the red dashed line shows the average for the initial 5 minutes. (MAKE SURE? Seems off) **B**, Spontaneous AP frequency in control and following the application of CBD for PV-INs. **C**, Current-clamp recordings showing the membrane potential of SST-INs during the application of CBD. **D**, Spontaneous firing rate of SST-INs before and after CBD application. Data points show mean ± SEM.

**Supplementary Figure 5.**
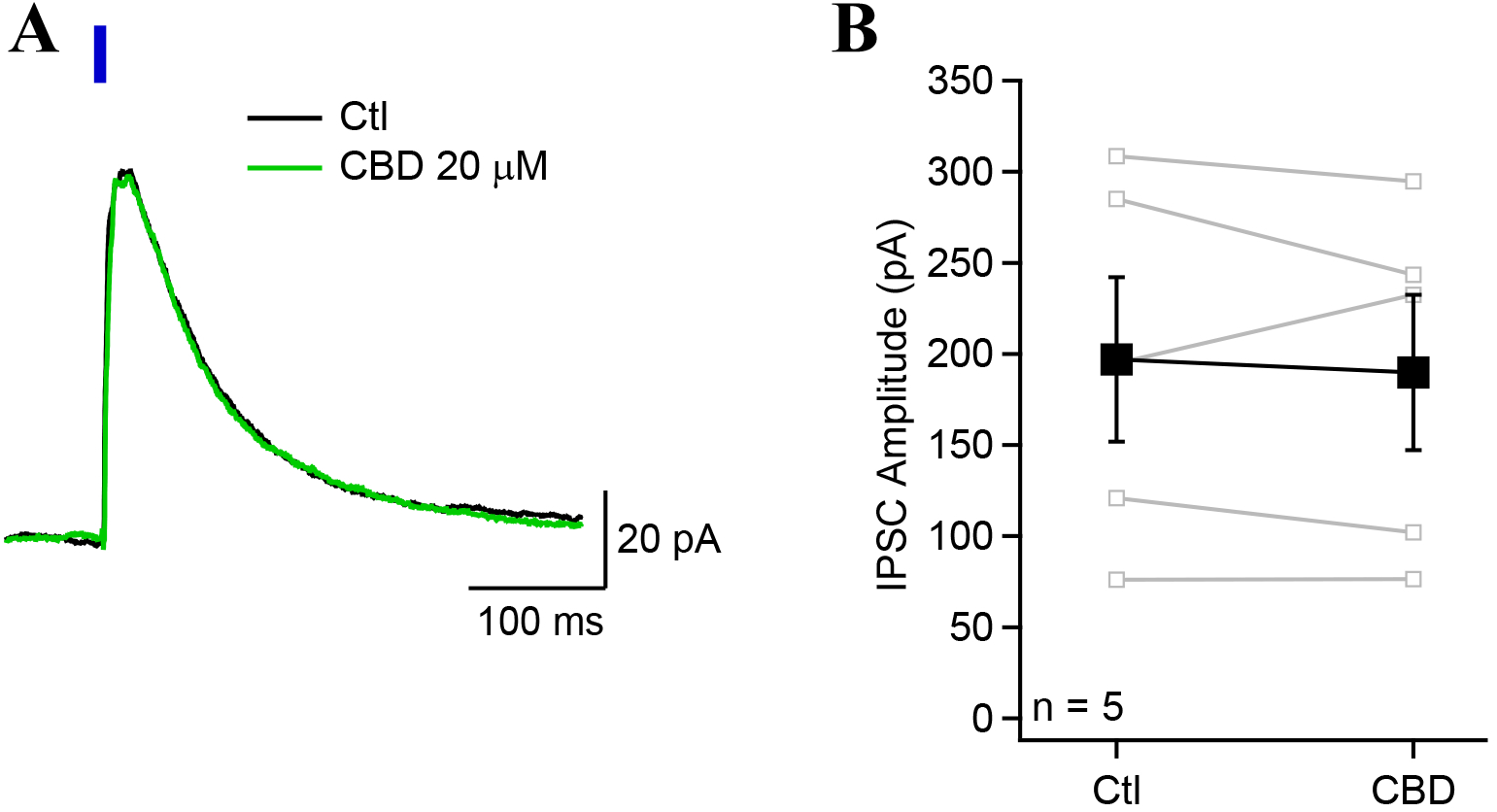
Cannabidiol does not affect the amplitude of IPSCs evoked from PV-INs. **A**, Voltage-clamp recordings (0 mV) from a CA1-PYR showing an IPSC evoked by blue light illumination in a slice from PV-Ai32 animal. Application of CBD had no effect on the IPSC amplitude (green trace). **B**, Optogenetically-evoked monosynaptic IPSC amplitude was not affected by the application of CBD.

**Supplementary Figure 6.**
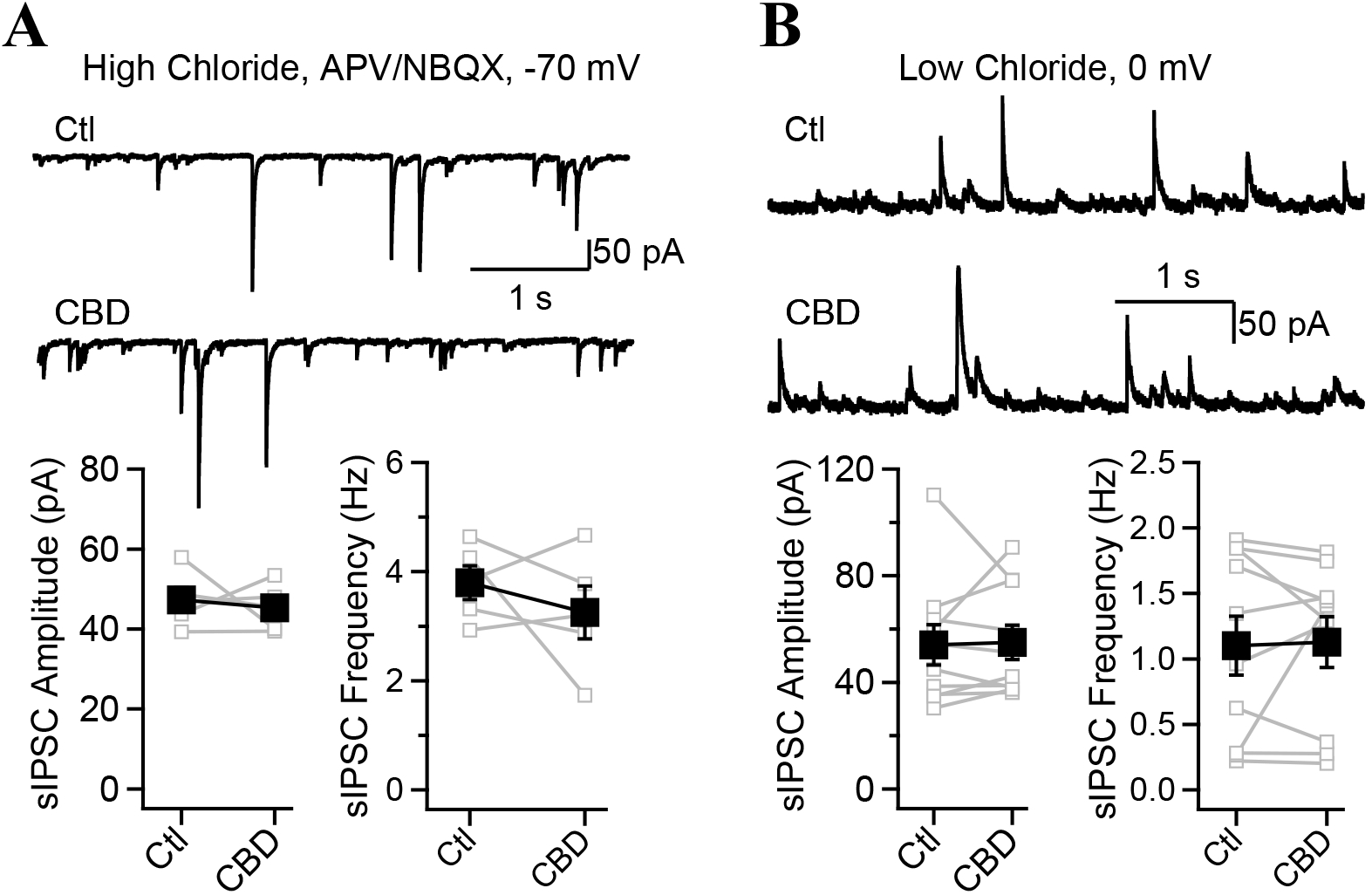
Cannabidiol does not affect spontaneous IPSCs recorded in CA1-PYRs. **A**, Spontaneous IPSCs recorded in voltage-clamp at –70 mV with a high chloride internal solution and in presence of NBQX and APV to isolate large inward inhibitory currents. Summary graphs show that CBD had no effect on the amplitude or the frequency of sIPSCs. **B**, Spontaneous IPSCs recorded with a low chloride intracellular from a CA1-PYR voltage-clamped at 0 mV before and after the application of CBD. The amplitude and the frequency of sIPSCs were unaffected by CBD. Data points show mean ± SEM.

**Supplementary Figure 7.**
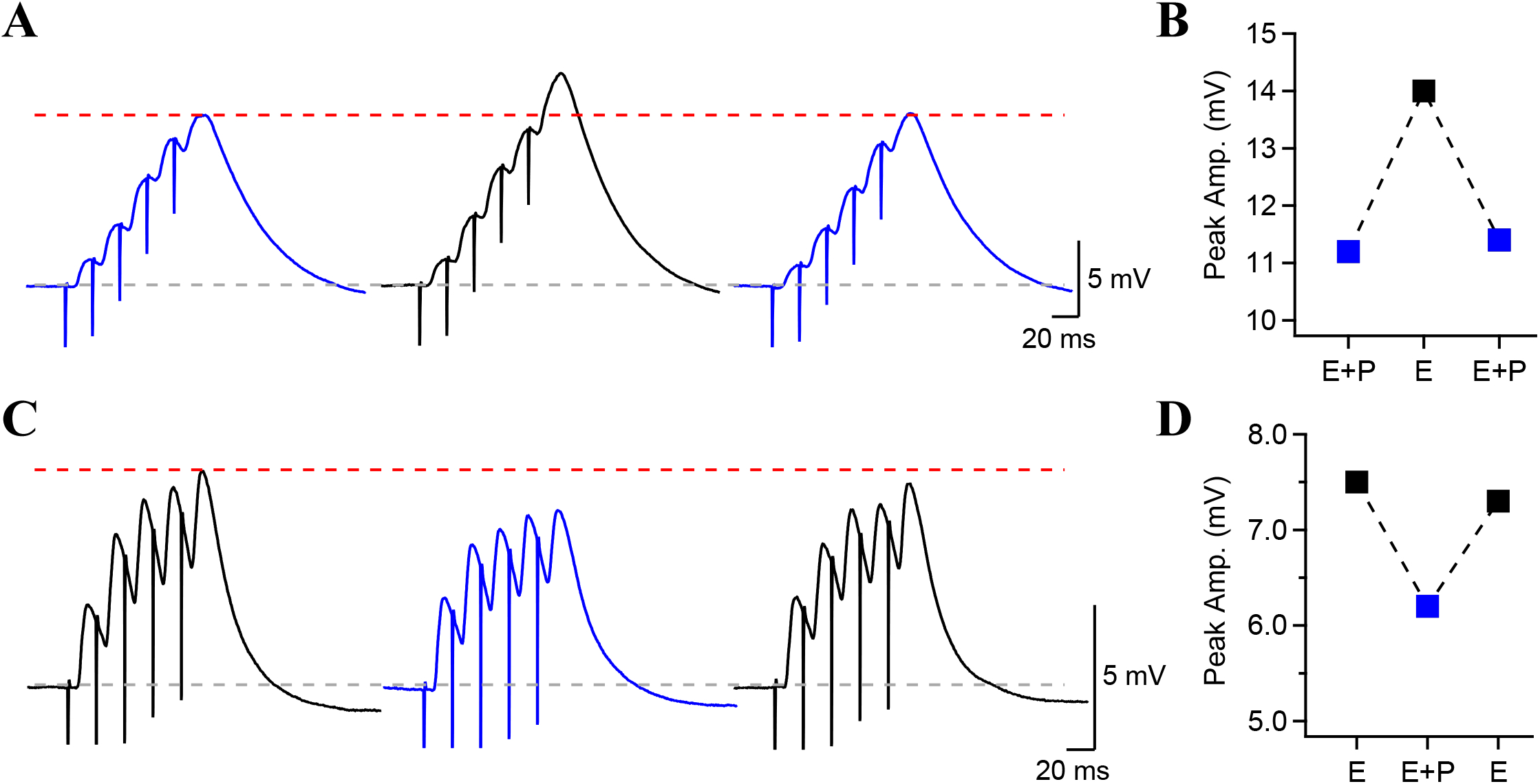
Reproducibility and reversibility of optogenetic PV-INs stimulation. **A**, Synaptic potentials evoked together with blue light (blue traces) or without (black trace). Recording were 3 minutes each, presented in the order of acquisition. **B**, Peak synaptic potential amplitude for electrical stimulation alone or combined electrical and optical stimulation. Photostimulation effect was reproducible. **C**, Synaptic potentials evoked with electrical stimulation with and without optogenetics stimulation. **D**, Peak synaptic potential amplitude during electrical and combined electrical and photostimulation. The effect of PV+ terminals photostimulation is reversible.

**Supplementary Figure 8.**
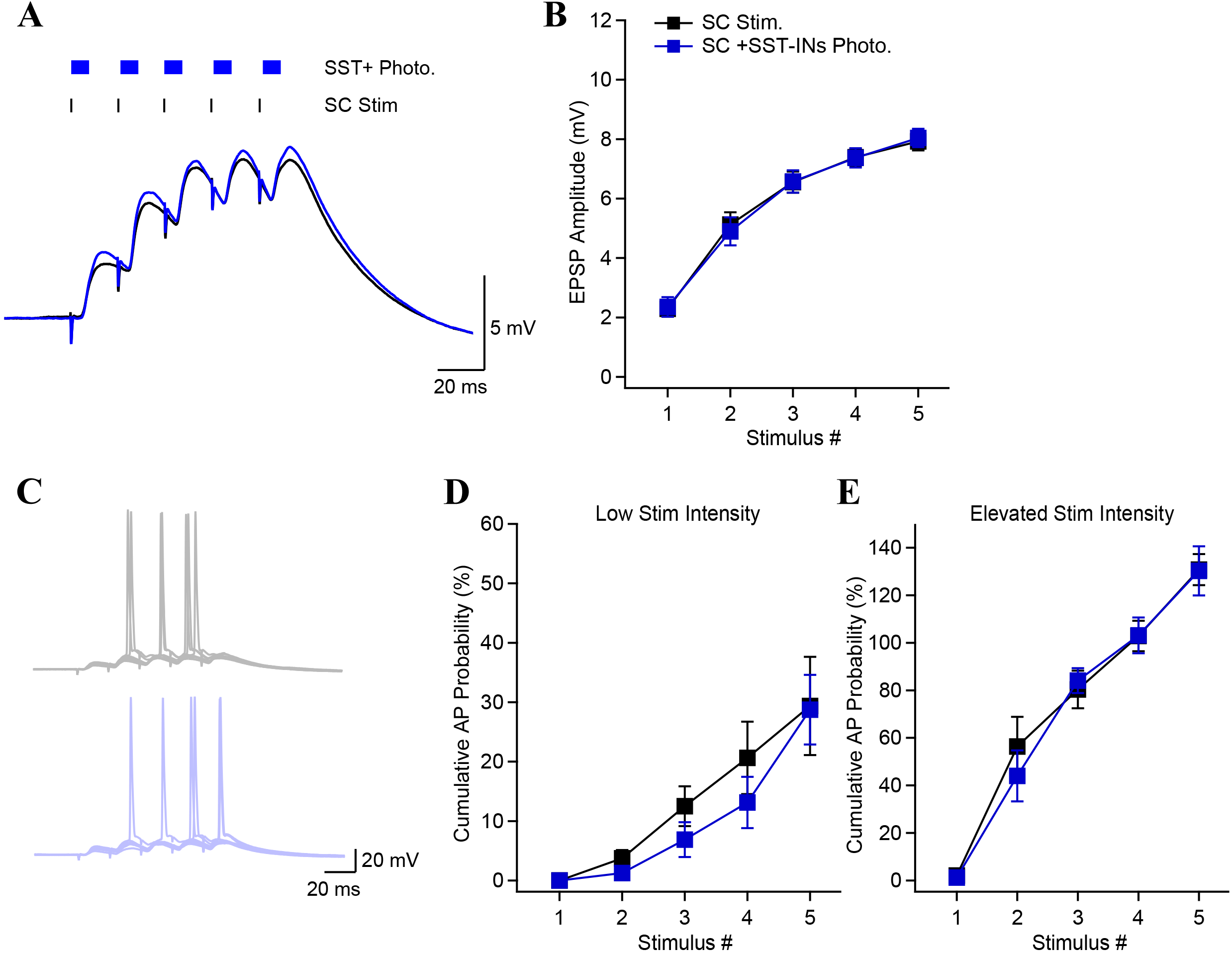
On demand SST-INs recruitment does not affect synaptic potential amplitude. **A**, Synaptic potentials evoked by SC stimulation, with and without concomitant optogenetic activation of SST-INs. **B**, EPSP amplitude as a function of stimulus number in control condition and during photostimulation of SST-INs. Unlike PV-INs photostimulation, SST-INs photoactivation did not decrease the EPSP amplitudes. **C**, Action potentials recorded from a CA1-PYR evoked by SC stimulation (10 consecutive sweeps) in absence (black) and with photostimulation (blue). **D**, **E**, Photostimulation of SST-INs had no effect on the cumulative AP probability during brief burst stimulation. This was observed under conditions of low (**D**) and high (**E**) stimulation intensity.

## Author contributions

SC, ECR, OD and RWT designed research. SC, ECR, ERN performed research and analyzed data. SC and RWT wrote the manuscript with input from all authors.

## Acknowledgements

This work was supported by grants to R.W.T. from the NIMH (MH071739), NIDA (DA040484), the Simons Foundation, and the Vulnerable Brain Project. S.C. was supported by a Charles H. Revson Senior Fellowship in Biomedical Science, Andrew Ellis and Emily Segal Investigator Grant from the Brain and Behavior Research Foundation (NARSAD), a postdoctoral fellowship from the Fonds de Recherche du Québec – Santé, and a K99/R00 Pathway to Independence Award from NIMH (K99MH126157). S.C. and O.D. received funding from Finding A Cure for Epilepsy and Seizures (FACES). E.R. was supported by the Ruth L. Kirschstein National Research Service Award (F30NS100293) and the NYU MSTP training grant (T32GM007308).

